# Reprogramming the Immune Suppressive Tumor Microenvironment in Glioma Enhances the Efficacy of Immune-Mediated Gene Therapy

**DOI:** 10.1101/2025.11.11.687828

**Authors:** Brandon L. McClellan, Jorge A. Peña Agudelo, Anzar A. Mujeeb, Ali A. Dabaja, Ziwen Zhu, Sadhakshi Raghuram, Maria Luisa Varela, Claire Tronrud, Kaushik Banerjee, Abraham Wei, Cecilia Calatroni, Lucia H. Zhang, Lissa Cruz Romero, Paul Oh, Mahmoud S. Alghamri, Andrew Robbins, Matthew D. Perricone, Ying Wang, Brian Shay, Peter Sajjakulnukit, Costas A. Lyssiotis, Joshua D. Welch, Anna Schwendeman, Pedro R. Lowenstein, Maria G. Castro

## Abstract

Gliomas account for ~80% of primary malignant brain tumors. Many CNS WHO grade 2–3 and some grade 4 gliomas harbor mutant isocitrate dehydrogenase 1 (mIDH1), which causes a gain of function mutation (IDH1 R132H) leading to the production of 2-hydroxyglutarate (2HG). Mutant IDH1-induced 2HG, through epigenetic reprogramming elicits an immune-permissive tumor microenvironment (TME). An immunosuppressive mechanism in the glioma TME involves adenosine production via the ectoenzyme CD73. This study investigates mIDH1’s influence on CD73 expression and adenosine levels. We demonstrate that mIDH1 glioma cells exhibit reduced CD73 expression, driven by DNA hypermethylation, leading to reduced adenosine levels. Since wtIDH1 gliomas have high CD73 expression, we evaluated CD73 blockade as an immunotherapy target. We show that CD73 inhibition used as monotherapy, did not improve survival in wtIDH1 glioma-bearing mice. However, when combined with immune-stimulatory Ad-TK (adenoviral vectors encoding herpes simplex virus thymidine kinase) and Ad-Flt3L (adenoviral vectors encoding FMS-like tyrosine kinase 3 ligand) gene therapy, CD73 blockade significantly enhanced therapeutic efficacy and increased anti-glioma effector T cell activity. These findings reveal that CD73 inhibition used in combination with immune stimulatory Ad-TK/Ad-Flt3L gene therapy may be an effective treatment for wtIDH1 gliomas, which could be readily translated to the clinical arena.

## Introduction

Gliomas are tumors of the central nervous system (CNS), accounting for roughly 26% of all CNS tumors^1,2^. The current standard of care (SOC) includes maximal surgical resection followed by radiotherapy and temozolomide chemotherapy. Despite the SOC, survival remains dismal with the median survival ranging from 8 months to 16.6 years, depending on the glioma type^1,2^. Gliomas are stratified into types based on several features which affect clinical outcomes. These features include genetic and epigenetic signatures of the glioma cells^3^. One such defining signature is the mutation status of isocitrate dehydrogenase gene 1 (IDH1)^3^. The most prevalent mutation of IDH1 is the replacement of arginine (R) to histidine (H) at amino acid residue 132 (R132H)^4,5^. Mutant IDH1-R132H (mIDH1) is a gain-of-function mutation resulting in the conversion of alpha-ketoglutarate (αKG) to the oncometabolite D-2-hydroxyglutarate (2HG)^6^. 2HG behaves as a competitive inhibitor of αKG dependent histone and DNA demethylases, including ten-eleven translocation (TET) hydroxylases and the Jumonji-C domain containing histone demethylases^7^. This widespread inhibition leads to a global hypermethylation phenotype^7^. A key feature of this hypermethylation is the aberrant methylation of CpG islands, especially within promoter regions, which disrupts normal gene expression programs^8^. These changes profoundly reshape glioma biology, affecting cell fate decisions, metabolic processes, genomic integrity, and interactions within the glioma tumor microenvironment (TME)^9–13^.

With relatively high numbers of immune suppressive cells, e.g. tumor associated macrophages (TAMs) and myeloid-derived suppressor cells (MDSCs), and high expression of inhibitory molecules, such as immune checkpoint proteins, the glioma TME features a strong anti-inflammatory landscape driven by multiple immune regulatory mechanisms^14^. Research has highlighted that mIDH1 has a significant impact on the glioma tumor immune response leading to an immune permissive TME^10,11,15^. Mutant IDH1 has been shown to alter the glioma TME by causing the generation of non-inhibitory neutrophils in lieu of polymorphonuclear-myeloid-derived suppressor cells (PMN-MDSCs)^11^. Mutant IDH1 glioma cells, when compared to wildtype IDH1 (wtIDH1) glioma cells, also exhibit hypermethylation of the *CD274* gene promoter region resulting in lower expression levels of immunosuppressive ligand PD-L1^10^. Mutant IDH1-mediated immune response differences, such as the ones mentioned above, play an important role in tumor progression and affect potential targets of immunotherapies^10,11^. Although several mIDH1-induced glioma immune response differences have been elucidated, many mechanisms of glioma TME immune suppression remain uninvestigated. Addressing this knowledge gap may uncover additional targets for precision therapies for gliomas based on IDH1 mutation status.

Along with other immune inhibitory pathways, adenosine signaling is a mechanism that allows gliomas to evade immune surveillance by inhibiting anti-tumor cytotoxic T cells^16^. The adenosinergic pathway converts the inflammatory metabolite ATP into the anti-inflammatory metabolite adenosine via ecto-enzymes CD39 and CD73^16,17^. Emerging evidence suggests that mIDH1 may modulate adenosinergic pathway signaling by influencing the expression of CD73^18^. *Braganhol et al.* used in silico analysis transcriptomic analysis and found that wtIDH1 glioblastomas exhibit significantly higher *NT5E*, CD73-encoding gene, expression compared with mIDH1 glioblastomas^18^. Since the adenosinergic pathway has been identified as a dominant axis of immunosuppression in gliomas and a key target for potential immunotherapies^16,19^, we sought to uncover the impact of mIDH1 on CD73 expression in gliomas and investigate CD73’s potential as an immunotherapy target.

Here, we unraveled a novel mechanism by which mIDH1 reshapes the glioma TME through the epigenetic suppression of CD73 expression, thereby leading to a reduction in extracellular adenosine—an immunosuppressive metabolite. We demonstrate that while CD73 inhibition alone is insufficient to impact wtIDH1 glioma survival, its combination with immune-stimulating Ad-TK/Ad-Flt3L (adenoviral vectors encoding herpes simplex virus 1–thymidine kinase/Feline McDonough sarcoma (Fms)–like tyrosine kinase 3 ligand) gene therapy significantly prolongs median survival and enhances anti-tumor immunity. These findings collectively highlight the distinct immune landscape conferred by the IDH1 mutation and validate the therapeutic potential of a combined CD73 blockade with Ad-TK/Ad-Flt3L gene therapy strategy to overcome the profound immune suppression characteristic of wtIDH1 gliomas, offering a promising avenue for clinical translation.

## Results

### Glioma Cells Harboring mIDH1 Have Decreased CD73 Expression

We analyzed The Cancer Genome Atlas (TCGA) datasets to assess the expression of adenosinergic pathway genes in the context of mIDH1 gliomas. Stratifying the TCGA glioblastoma low-grade glioma (GBMLGG) cohort patient data into wtIDH1 and mIDH1 groups revealed that CD73-encoding gene, *NT5E*, expression was significantly downregulated in patients with mIDH1 (Figure 1A). Because mIDH1 is commonly found in both glioma and acute myeloid leukemia (AML)^9^, the TCGA-AML cohort was also interrogated for mIDH1 differences. Similarly to glioma, *NT5E* mRNA expression was also decreased in mIDH1 AML patients compared to those of wtIDH1 AML patients (Figure S1).

**Figure 1:**
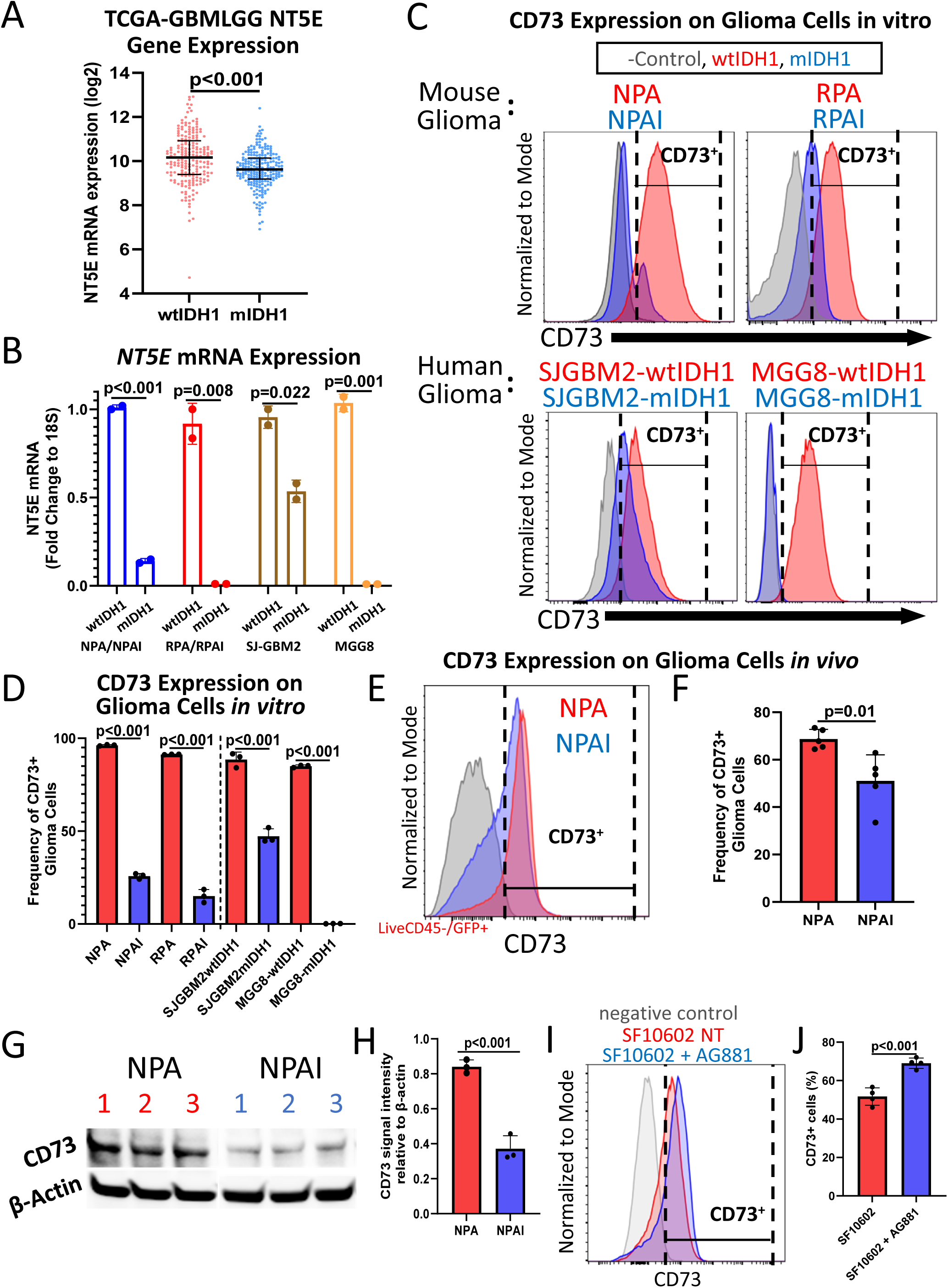
Mutant-IDH1 Glioma Cells Have Decreased CD73 Expression. (A) Expression of *NT5E* mRNA based on TCGA-GBM and TCGA-LGG combined dataset stratified based on the wildtype (*n* = 204) and mutant (*n* = 230) IDH1 status of the patients (Mann–Whitney U test). (B) Quantitative PCR analysis of *NT5E* mRNA expression across wtIDH1 glioma cells (NPA, RPA, SJGBM2-wtIDH1, MGG8-wtIDH1) and mIDH1 glioma cells (NPAI, RPAI, SJGBM2-mIDH1, MGG8-mIDH1) (Student’s t test, *n* = 2). (C) Representative flow cytometry histograms and (D) quantitation of the percentage of wildtype and mutant IDH1 glioma cells expressing CD73. (Student’s t test, *n* = 3). (E) Representative flow cytometry histogram and (F) quantitation of the frequency of CD73 expressing glioma cells to total glioma cells. (Student’s t test, *n* = 5). (G) Western blot image and (H) quantitation of CD73 levels from total glioma tumor in vivo (Student’s t test, *n* = 3). (I) Representative flow cytometry histogram and (J) quantitation of CD73 expressing human mIDH1 glioma cells (SF10602s) with and without mIDH1 inhibitor (AG881) treatment (Student’s t test, *n* = 4).

Tumor cells have been identified as the primary source of CD73 in gliomas^20^, so we investigated the CD73 difference by measuring glioma cell CD73 expression. CD73 expression on the cell surface of wtIDH1 glioma cell lines (NPA, RPA, SJ-GBM2-wtIDH1, MGG8-wtIDH1) were compared to their syngeneic mIDH1 counterparts (NPAI, RPAI, SJ-GBM2-mtIDH1, MGG8-mIDH1) using reverse transcription quantitative polymerase chain reaction (RT-qPCR). Mouse glioma cells (NPA, NPAI, RPA, RPAI) were generated using the Sleeping Beauty transposon system to insert shP53 and shATRX with or without mIDH1-R132H^11,21,22^. The shP53 and shATRX constructs mimic the loss-of-function mutations in TP53 and ATRX commonly found in astrocytomas, which frequently co-occur with mIDH1^3,23^. SJ-GBM2-wtIDH1, SJ-GBM2-mIDH1, MGG8-wtIDH1, MGG8-mIDH1 are patient-derived glioma cells^11^. The RT-qPCR results demonstrated that mIDH1 glioma cells have decreased *NT5E* transcription levels (Figure 1B). To assess the glioma cell CD73 protein levels, we measured the surface CD73 levels. Consistent with the transcript data, mIDH1 glioma cells exhibited decreased surface expression of CD73 compared to wtIDH1 cells (Figures 1C and 1D).

To measure the CD73 expression levels on the surface of wtIDH1 and mIDH1 glioma cells *in vivo*, we intracranially implanted NPA (wtIDH1) and NPAI (mIDH1) glioma cells, which are GFP+, to generate immune-competent glioma mouse models (GMMs). Upon reaching late-stage symptomatic phenotype, we collected the tumor tissue and measured glioma cell (Live+CD45-GFP+) CD73 expression. From the GMM TME, the mIDH1 glioma cells maintained decreased CD73 in comparison to wtIDH1 glioma cells (Figures 1E and 1F). We then isolated the late-stage tumors from GMMs and measured the total CD73 expression. The results show that mIDH1 gliomas tumors have less overall CD73 compared to wtIDH1 gliomas (Figures 1G and 1H).

We validated if the reduction of CD73 is attributable to the mutation of IDH1 using primary human mIDH1 glioma cells, SF10602s^24^. SF10602 cells were treated with vorasidenib (AG881) to potently inhibit mIDH1 and examine the effect on CD73^22,25^. Consistent with the prior results, mIDH1 inhibitor treatment led to an increase in CD73 expression by SF10602 cells (Figures 1I and 1J), resembling that of wtIDH1 glioma cells. These findings provide clear evidence that mIDH1 causes a decrease in CD73 expression by glioma cells.

### Mutant IDH1 Leads to Epigenetic Downregulation of CD73 Expression by Glioma Cells

The mutation of IDH1 alters the glioma cells by generating the oncometabolite 2HG, which then inhibits DNA hydroxylases and histone demethylases^26^. In particular, 2HG reduces the conversion of 5-methylcytosine (5mC) to 5-hydroxymethylcytosine (5hmC) by inhibiting TET2^26^. This causes the hypermethylated DNA phenotype known as glioma CpG island methylator phenotype (G-CIMP)^8,26^. Since this hypermethylated CpG island state is genome wide, we hypothesize that the reduction of CD73 is due to mIDH1-induced epigenetic reprogramming. To test this hypothesis, we conducted enzyme-based methylation sequencing (methyl-seq) and examined the methylation around the promoter region of *NT5E*^27^. Wildtype IDH1 and mIDH1 glioma cells’ DNA was preprocessed, 5mC’s were oxidized to 5hmCs, and the unmodified cytosines were deaminated to uracils before amplification and sequencing (Figure 2A)^27^. Whole genome analysis revealed that mIDH1 glioma cells’ *NT5E* promoter region had increased methylation at the CpG islands (Figure 2B). Increased DNA methylation corresponds to lower transcription^28^, thereby indicating that this increased methylation may be responsible, at least in part, for the decrease in CD73 expression.

**Figure 2:**
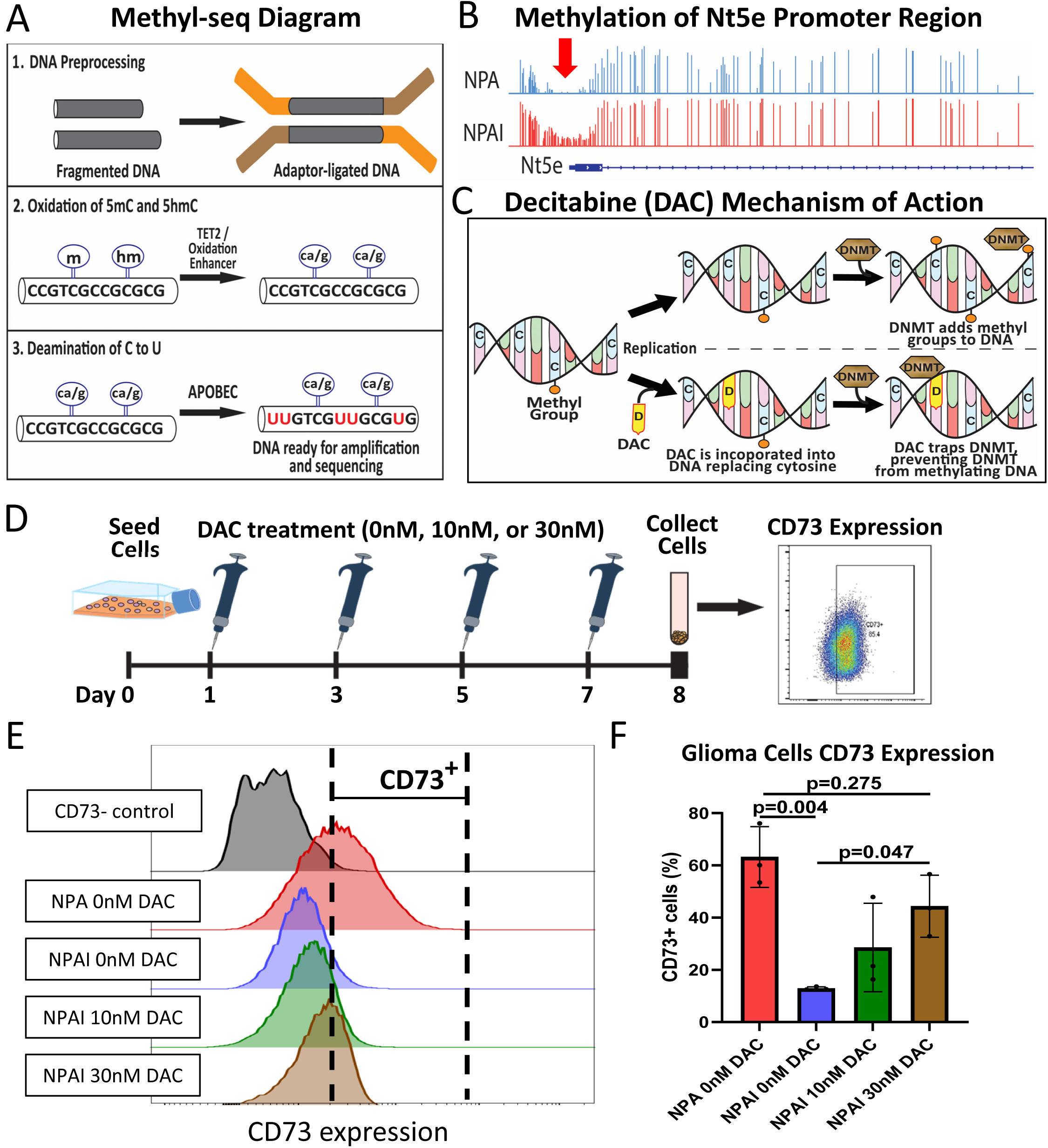
CD73 Expression Is Epigenetically Downregulated in mIDH1 Glioma Cells. (A) Mechanism of methylation sequencing (methyl-seq) DNA methylation profiling. (B) Methyl-seq readout of the CD73 encoding gene region for NPA (wtIDH1) and NPAI (mIDH1). Red arrow indicates the *NT5E* promoter region with notable methylation difference between NPA and NPAI glioma cell DNA methylation. (C) Decitabine (DAC) mechanisms of action resulting in DNA methylation by inhibiting DNMT. (D) Timeline of DAC treatment on glioma cells to measure the effect on CD73 expression. (E) Representative flow cytometry histograms and (F) quantitation of CD73 expression from wtIDH1 and mIDH1 glioma cells treated with DAC (one-way ANOVA, *n* = 3).

We sought to validate the connection between DNA methylation and CD73 expression by treating wtIDH1 and mIDH1 glioma cells with decitabine (5-aza-2′-deoxycytidine; DAC). DAC is an FDA-approved hypomethylating agent that can incorporate into replicating DNA, where it irreversibly binds and inactivates DNMT1. This results in inhibition of DNMT-mediated DNA methylation (Figure 2C), leading to a reduction in the mIDH1-induced DNA hypermethylation. Treatment of glioma cells with DAC was carried out over one week to allow the epigenetic changes to take effect (Figure 2D). DAC treatment of mIDH1 NPAI cells increased CD73 expression from the NPAI cells’ decreased baseline level to a level not significantly different from that of wtIDH1 NPA cells (Figures 2E and 2F). Together, these data indicate that the mutation of IDH1 causes decreased CD73 expression due to increased glioma cell DNA methylation, particularly at the *NT5E* promoter region.

### Mutant IDH1 Glioma Cell Conditioned Media Has Reduced Adenosine Levels

The glioma TME is marked by high levels of cell damage, hypoxia, and cell death^14,29^. These characteristics contribute to high levels of extracellular ATP (eATP)^30,31^. Although eATP is an immune-stimulatory metabolite, the glioma TME converts it into the immunosuppressive metabolite adenosine through the adenosinergic pathway (Figure 3A). The initial step in this pathway is the conversion of eATP to AMP via two sequential dephosphorylation reactions by CD39. Then, CD73 performs the final dephosphorylation reaction converting AMP to adenosine^16^. To begin assessing the effects of the mIDH1 induced CD73 reduction in glioma cells, we examined the enzymatic function of CD73. We measured the levels of the CD73 substrate and product, AMP and adenosine respectively, in the conditioned media of wtIDH1 and mIDH1 glioma cells. Results showed that mIDH1 glioma cells have increased concentration of AMP in their conditioned media compared to that of their syngeneic wtIDH1 glioma cells (Figures 3B and 3D). This AMP change was accompanied by a decrease in adenosine levels in the conditioned media (Figures 3E and 3G). These results substantiate that there is reduced CD73 enzymatic function of converting AMP to adenosine from the mIDH1 glioma cells, and this is likely due to the reduction of CD73 expression.

**Figure 3:**
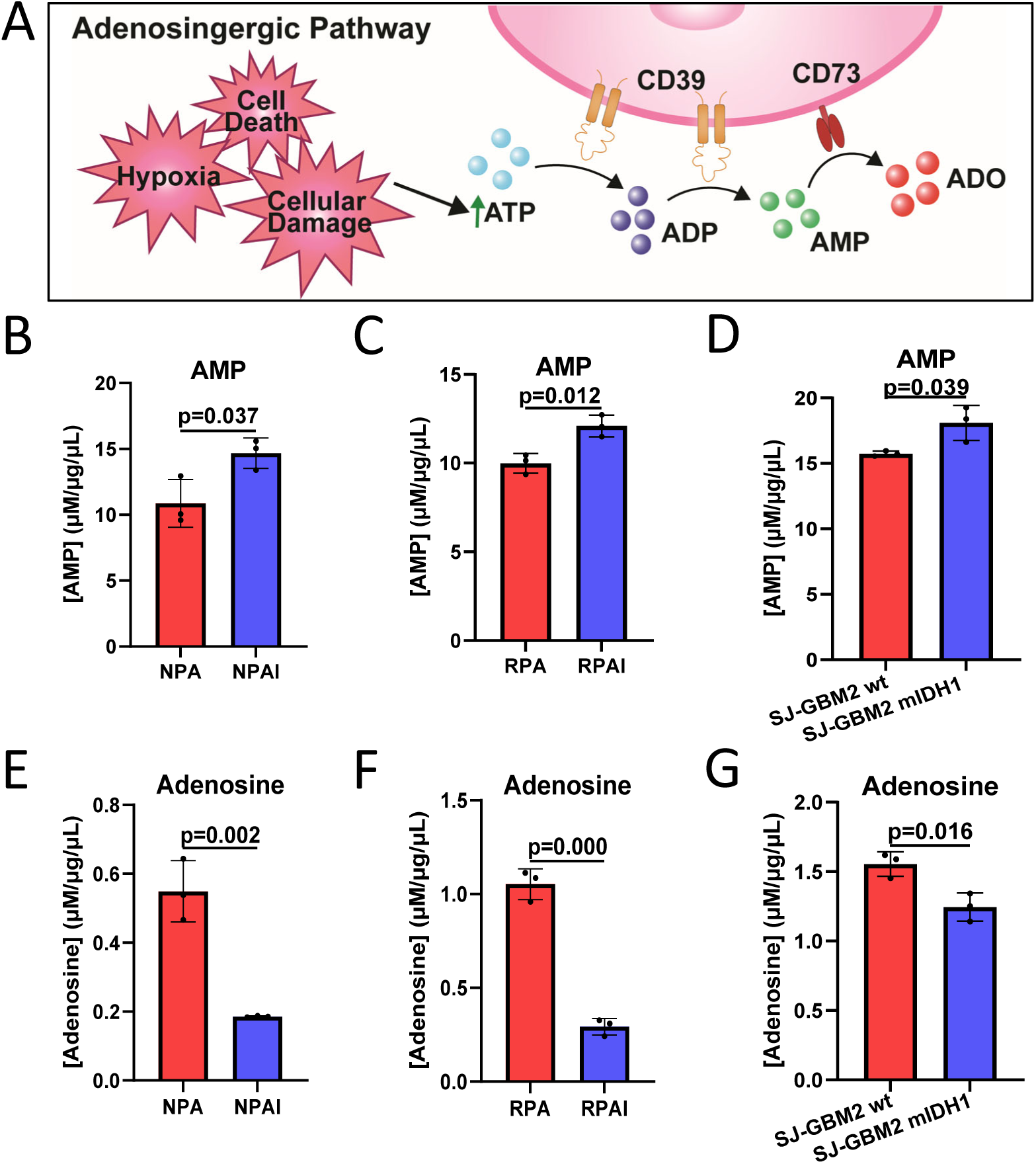
Adenosinergic Pathway Related Metabolites. (A) Adenosinergic pathway diagram. (B-D) AMP measurement of conditioned media, normalized to total protein, of wtIDH1 glioma cells (NPA, RPA, SJGBM-wt) compared to mIDH1 glioma cells (NPAI, RPAI, SJ-GBM2) (Student’s t test, *n* = 3). (E-G) Adenosine measurement of conditioned media, normalized to total protein, of wtIDH1 glioma cells compared to mIDH1 glioma cells (Student’s t test, *n* = 3).

### Glioma Cell CD73 Expression Has Minimal Effect on Glioma Survival

CD73 as a part of the adenosinergic pathway is one of the many pathways that contributes to glioma induced immunosuppression^16^. Since wtIDH1 gliomas were shown to have high CD73 expression, we investigated the effect of glioma cell CD73 expression on glioma survival by generating CD73 knockout (CD73KO) wtIDH1 GMMs. The Sleeping Beauty (SB) transposase system was employed to generate the CD73KO gliomas de novo^11,21^. Gliomas were induced in wildtype and *NT5E*^-/-^ mice by NRAS overexpression in combination with shP53 and shATRX (Figure 4A). Tumors developed in the wildtype mice (NRAS/shP53/shATRX; NPA CD73wt) and *NT5E*^-/-^ mice (NRAS/shP53/shATRX/*NT5E*^-/-^; NPA CD73KO). Regardless of CD73 presence, the SB GMMs had similar median survivals of 63.5 days with CD73 and 64.5 days without CD73 (Figure 4B). Using the combined glioma TCGA patient dataset, TCGA GBMLGG, we stratified patients’ survival based on high (top 25^th^ percentile) and low (bottom 25^th^ percentile) *NT5E* expression. Given that mIDH1 glioma patients typically have longer survival than wtIDH1 patients and *NT5E* expression is lower in mIDH1 cases, we removed the confounding effect of IDH status by stratifying the data further based on IDH1 mutation status^32^. After IDH1-based stratification, *NT5E* expression did not correlate with changes in patient survival within either IDH1 status subgroup (Figures 4C and 4D).

**Figure 4:**
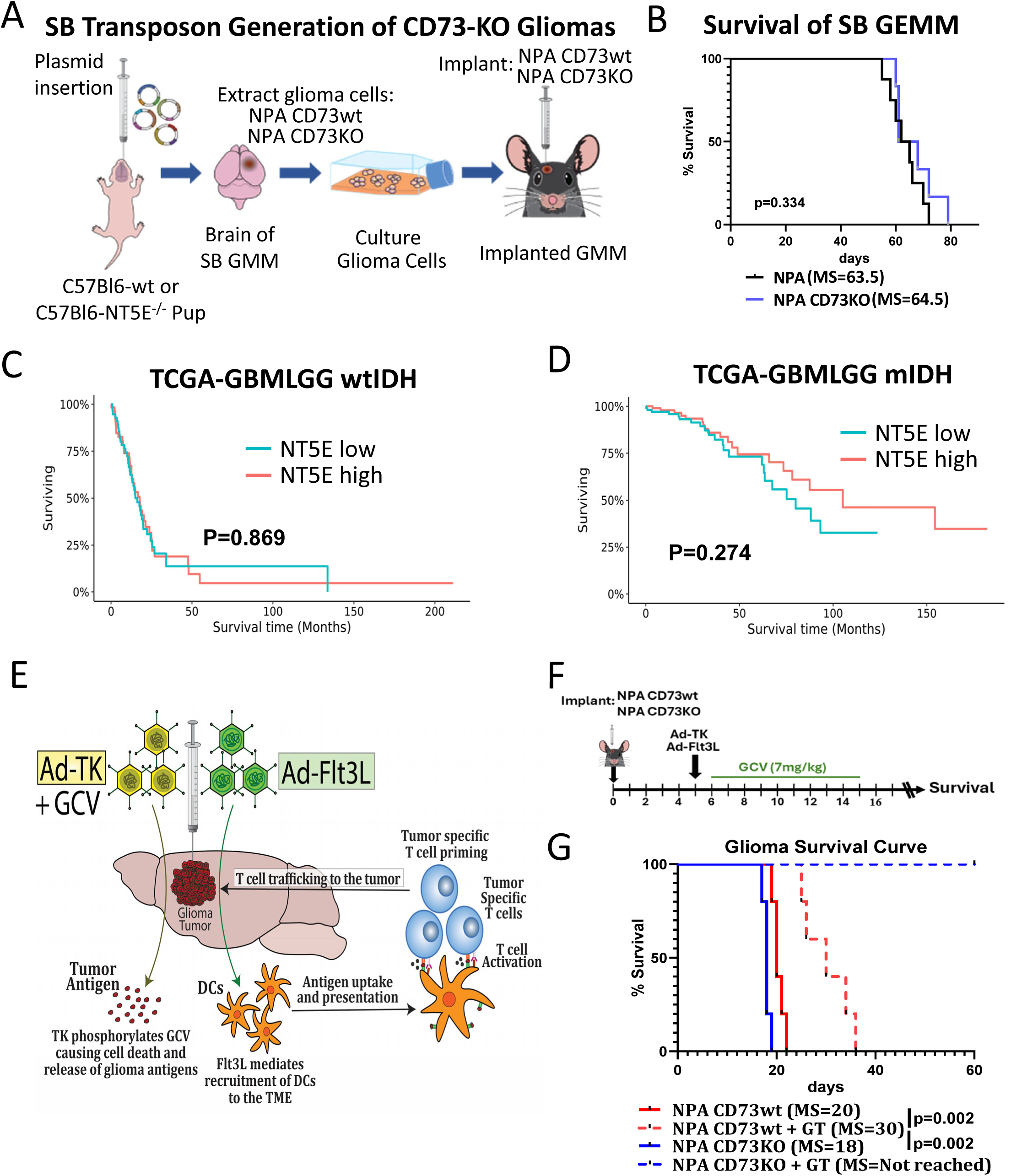
Role of Glioma Cell CD73 in Tumor Progression. (A) Diagram of Sleeping Beauty (SB) transposon generated CD73 knockout (NPA CD73KO) and CD73 wildtype (NPA CD73wt) glioma models and the generation of implanted glioma mouse models (GMMs). (B) Survival of SB generated CD73-wt glioma models (NPA; *n* = 8) compared to CD73-KO glioma models (NPA CD73KO; *n* = 6) (Log-rank test). (C) TCGA (GBMLGG) wtIDH patient survival curve with patient cases stratified on *NT5E* gene expression (top 25% cases are high *NT5E n* = 58; bottom 25% of cases are low *NT5E n* = 59) (Mann-Whitney U test). (D) TCGA (GBMLGG) mIDH patient survival curve with patient cases stratified on *NT5E* gene expression (top 25% cases are high *NT5E n* = 107; bottom 25% of cases are low *NT5E n* = 108) (Mann-Whitney U test). (E) Mechanism of action of Ad-TK/Ad-Flt3L gene therapy (GT). (F) Schematic showing experimental design of NPA CD73wt and NPA CD73KO glioma cell implantation and treatment with or without Ad-TK/Ad-Flt3L GT. (G) Survival curve of NPA CD73wt and NPA CD73KO glioma models (*n* = 5) treated on the timeline shown in (F).

The SB GMMs showed no difference in survival based on CD73 presence, yet these were total knockout models with CD73 absent on all cells. The lack of CD73 entirely may lead to extraneous variables affecting the survival, such as compensatory mechanisms. Therefore, since our goal was to evaluate the effect of glioma cell intrinsic CD73 on glioma survival, we cultured and implanted the NPA CD73wt and NPA CD73KO glioma cells from the SB GMMs into wildtype, immune competent mice (Figure 4A). The knockout of CD73 on the NPA CD73KOs was verified (Figures S2A and S2B). This generated GMMs with conditional knockouts of CD73 limited to the glioma cells.

Since CD73 did not affect survival in the SB GMMs (Figure 4B), we combined the CD73KO glioma cell implantation with Ad-TK/Ad-Flt3L gene therapy to determine if the lack of CD73 would support immune stimulating therapies. Ad-TK/Ad-Flt3L gene therapy works by a two-armed approach (Figure 4E)^33^. The Ad-TK encodes for herpes simplex virus type 1 thymidine kinase (TK), which activates ganciclovir (GCV) to cause immunogenic cell death. The Ad-Flt3L encodes for the cytokine Flt3L to recruit dendritic cells (DCs) to the glioma TME. These DCs capture tumor antigens from the Ad-TK induced-cell death mechanism, and stimulate tumor-specific T cells to generate a robust anti-tumor T cell response. GMMs were treated with Ad-TK and Ad-Flt3L on day 5 post-implantation, and GCV was administered from days 6-15 post-tumor cell implantation (Figure 4F). The glioma cell CD73KO alone did not alter survival in these models [median survival (MS): 18 days with CD73KO, MS: 20 days with CD73wt], but the absence of glioma cell CD73 extended the survival of the GMMs treated with Ad-TK/Ad-Flt3L gene therapy (MS: 30 days with gene therapy and CD73wt, MS: not reached with gene therapy and CD73KO) (Figure 4G). The CD73KO glioma models treated with Ad-TK/Ad-Flt3L gene therapy remained tumor free without further treatment (Figure 4G). Cumulatively, these data suggest that CD73 does not significantly influence glioma survival. Nevertheless, the findings offer insight into the potential role of CD73 as a target for immunotherapy, both in monotherapy and in combination with other immune-stimulating therapies.

### Targeting CD73 Boosts the Efficacy of Ad-TK/Ad-Flt3L Gene Therapy to Improve Survival

Based on effectiveness in extending the survival of Ad-TK/Ad-Flt3L gene therapy treated GMMs, we began examining CD73 as an immunotherapy target by inhibiting CD73 alone and in combination with the TK-Flt3L gene therapy. Combining these therapies may be necessary to extend glioma survival because without intervention, the glioma TME is highly immunosuppressive. While inhibiting CD73 alone reduces this immune suppression, the efficacy is limited by a scarce supply of anti-tumor immune cells. The addition of Ad-TK/Ad-Flt3L gene therapy addresses this limitation by increasing the number of DCs and cytotoxic CD8 T cells. Therefore, the simultaneous use of these two treatments is warranted: the GT generates the effector cells, and the CD73 inhibition reduces the immune barriers, allowing the increased CD8 T cells to successfully engage and attack the tumor (Figure 5A). Wildtype GMMs were treated with Ad-TK/Ad-Flt3L gene therapy, along with the small molecule inhibitor of CD73 (AB680) or the monoclonal antibody inhibitor of CD73 (aCD73) on the displayed timeline (Figure 5B). Anti-CD73 antibodies’ ability to cross the blood brain barrier and enter the brain was confirmed by probing mouse brain tissue sections with secondary antibodies against the host rat isotype Fc region of the antibodies (Figure S3). Treatment of the GMMs with CD73 inhibitors alone did not confer benefits as a monotherapy [no treatment MS: 20 days; aCD73 treated MS: 18 days; AB680 treated MS: 21 days] (Figure 5C). When CD73 inhibition was used in combination with the immune-stimulatory Ad-TK/Ad-Flt3L gene therapy (GT), CD73 inhibition resulted in extended survival. Median survival was 30 days for the GT treated group, which increased to 41 days for the combination treated GT aCD73 treated group, and 46 days for the combination GT AB680 treated group (Figure 5C). Notably, both combination treatments resulted in 20% long-term survivors (>80 days post glioma cell implantation).

**Figure 5:**
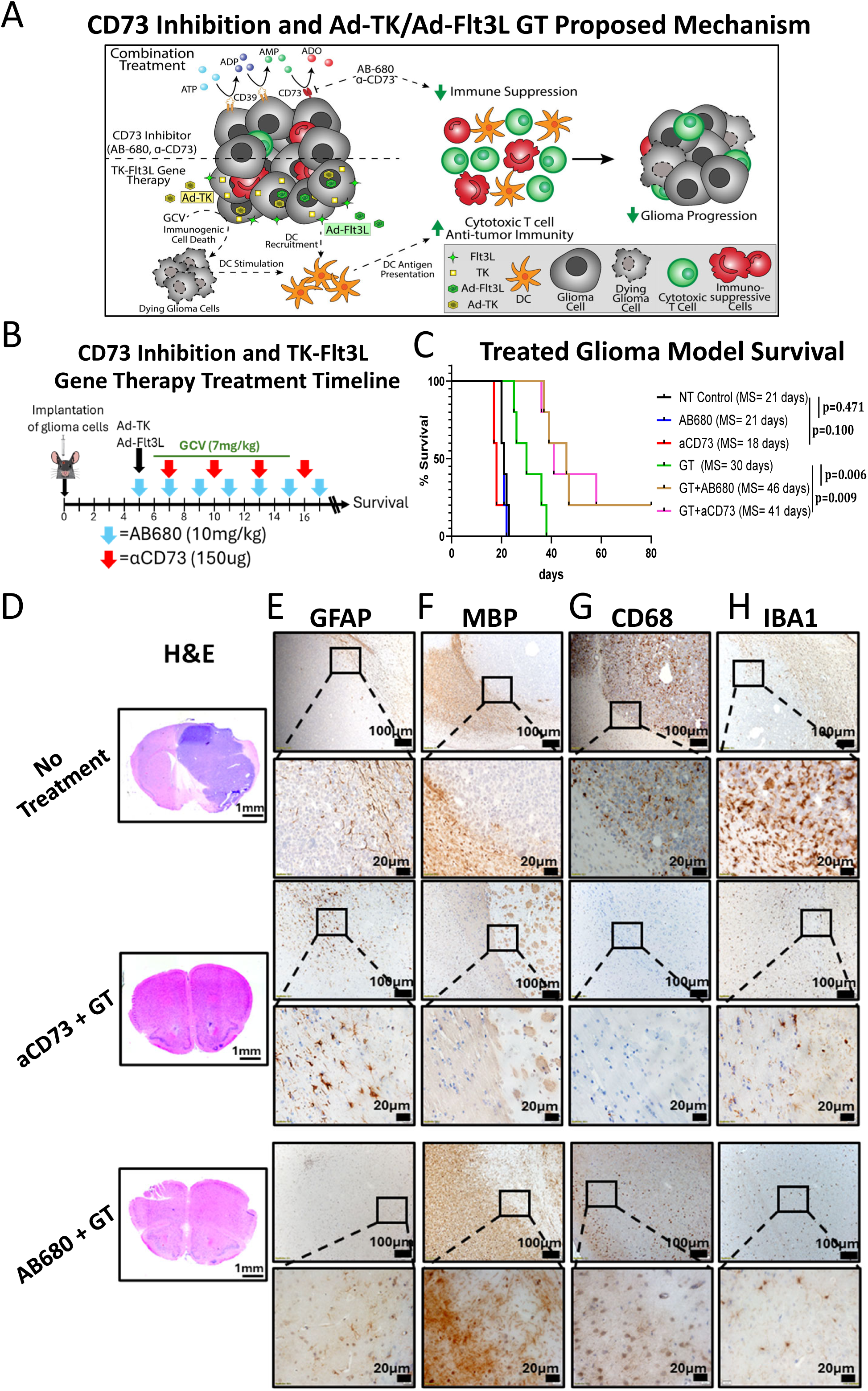
CD73 Inhibition as an Immunotherapy. (A) Proposed mechanism of action of the adenosinergic pathway mediated immune suppression contributing to glioma progression with the combination of CD73 inhibition with Ad-TK/Ad-Flt3L gene therapy. (B) Experiment timeline schematic of NPA wtIDH1 glioma cell implantation and treatments with CD73 inhibiting monoclonal antibody (aCD73), small molecule inhibitor of CD73 (AB680), and Ad-TK/Ad-Flt3L gene therapy. (C) Survival curve of NPA glioma models (*n* = 5) treated on the timeline shown in (B) (Log-rank test). (D) H&E staining of 5 µm paraffin-embedded brain sections from saline, long-term survivors from aCD73 GT and AB680 GT treatment groups (scale bar = 1 mm). Paraffin-embedded 5 µm brain sections for each treatment group were immunohistochemistry stained for (E) glial fibrillary acidic protein (GFAP), (F) myelin basic protein (MBP), (G) CD68, and (H) IBA1. Low magnification (10X) panels (black scale bar = 100 µm), high magnification (40X) panels (black scale bar = 20 µm) indicate positive staining for the areas delineated in the low-magnification panels. Representative images from a single experiment consisting of independent biological replicates are displayed.

H&E staining revealed the presence of tumors in the hemisphere region of the no treatment, AB680, GT alone, and aCD73 alone treated mice (Figures 5D and S4A). In contrast, microscopic examination showed no evidence of intracranial tumor in long-term survivors from combination aCD73 GT and AB680 GT treated groups (Figure 5D). No apparent areas of hemorrhages, necrosis, or invasion were present in the long-term survivors after receiving the combination therapies aCD73 GT and AB680 GT. To examine whether this combination treatment affected the surrounding brain architecture, we performed immunohistochemistry (IHC) staining using glial fibrillary acidic protein (GFAP) and myelin basic protein (MBP) as markers for myelin sheath integrity. Results showed no apparent changes in brain architecture in long-term surviving mice receiving the combined aCD73 GT and AB680 GT treatment (Figure 5E and 5F). While the no treatment, AB680, GT alone, and aCD73 groups show strong GFAP staining around the tumor and a marked loss of MBP signal in the peritumoral region (Figures 5E, 5F, S4B and S4C).

To further evaluate the level of immune cellular infiltrates, brain sections of mice from the experimental groups were analyzed by IHC using markers for macrophages (CD68) and microglia (IBA1). Visual inspection revealed increased infiltration of macrophages (CD68+ cells) within the TME and the adjacent brain parenchyma in the groups that received no treatment, AB680, GT alone, and aCD73 (Figures 5G and S4D). In contrast, the long-term survivors from combination aCD73 GT and AB680 GT treated groups show a reduced number of CD68+ macrophages in comparison to the other groups (Figure 5G). We also observed IBA1+ microglia surrounding brain parenchyma in the groups treated with saline, AB680, GT alone, and aCD73, whereas long-term survivors from combination aCD73 GT and AB680 GT treated groups show reduced levels of IBA1 expression (Figures 5H and S4E). Serum chemistry analysis and histopathological assessment of liver sections indicate no toxicity induced by the CD73 inhibitors and Ad-TK/Ad-Flt3L gene therapy treatments (Figures S5 and S6).

### CD73 Inhibition Improves CD8 T cells Anti-Tumoral Effector Functions in Combination with Ad-TK/Ad-Flt3L Gene Therapy

To provide a mechanistic basis for the improved survival observed in response to the Ad-TK/Ad-Flt3L GT and CD73 inhibition therapies, we examined their effect on the CD8 T cell population.

We generated GMMs using wtIDH1 glioma cells (NPA OVA cells). The GMMs were treated with CD73 inhibitor AB680 and Ad-TK/Ad-Flt3L GT. These mice were then euthanized 10 days post treatment (Figure 6A). Interferon gamma (IFNγ) and granzyme B (GzB) were measured as key effector molecules of anti-tumor CD8 T cells. The combination treated group had a higher frequency of CD8 T cells expressing IFNγ and GzB (Figures 6B and 6C). This suggests that compared to the monotherapies, the combination of Ad-TK/Ad-Flt3L GT and CD73 inhibition increases the antitumoral function of CD8 T cells, which would contribute to extending glioma survival.

**Figure 6:**
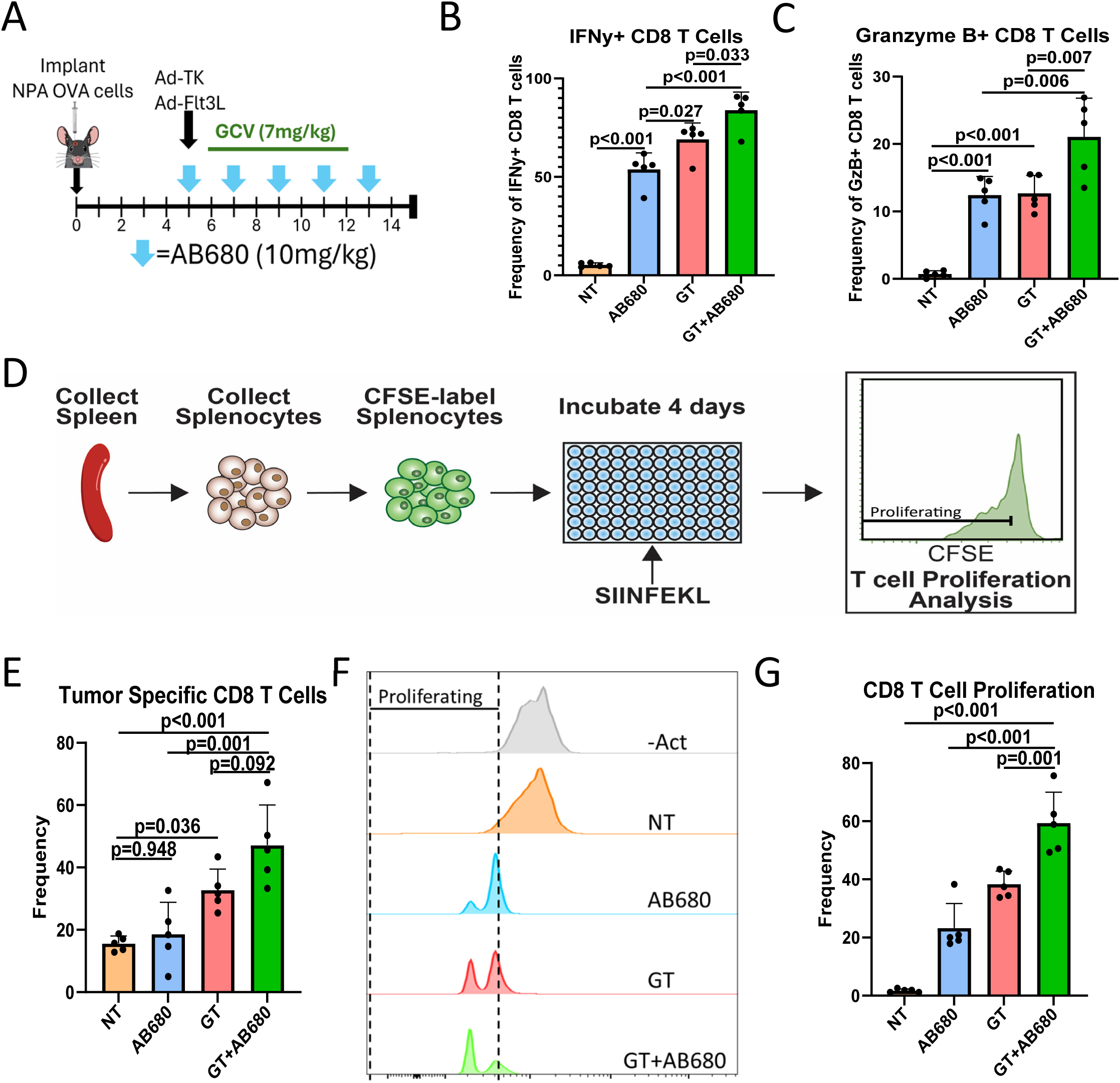
CD73 Inhibition and Ad-TK/Ad-Flt3L Gene Therapy Combination Induces Greater Effector Functions of Tumor Specific CD8 T cells. (A) Schematic showing experimental design of NPA glioma cell implantation and treatment with small molecule inhibitor of CD73 (AB680) and Ad-TK/Ad-Flt3L gene therapy. (B-C) Quantitation of the frequency of CD8 that are (B) IFNγ positive, (C) granzyme-B (GzB) positive (one-way ANOVA, *n* = 5). (D) Schematic showing experimental design of tumor specific CD8 T cell splenocyte activation and proliferation assay. (E) Flow cytometry quantitation data of tumor specific CD8 T cells from the splenocyte T cell proliferation experiment (one-way ANOVA, *n* = 5). (F) Representative flow cytometry histograms and (G) quantitation of CFSE expression as an indication of CD8 T cell proliferation (one-way ANOVA, *n* = 5).

To further validate the findings that the combination treatment stimulated anti-tumor CD8 T cell function, we examined the CD8 T cell proliferation capacity from splenocytes extracted from the treated GMMs implanted with NPA OVA cells and euthanized 10 days post treatment (Figure 6A). The splenocytes were isolated, stained with carboxyfluorescein diacetate succinimidyl ester (CFSE), and incubated with SIINFEKL for 4 days (Figure 6D). We then measured the frequency of surrogate tumor antigen (OVA) specific CD8 T cells and found that there were increased levels in the GT and AB680 GT combination treated groups compared to the no treatment control group (Figure 6E). There was, however, no significant difference with the addition of CD73 inhibition to the gene therapy (p=0.092, Figure 6E). In examining the CSFE dilution as a marker of proliferation, we found that the percentage of CD8 T cells proliferation for the untreated, AB680, GT, and AB680 GT combination treated groups were ~2%, ~23%, ~38%, and ~59% respectively (Figures 6F and 6G). The increased TME-infiltrating CD8 T cells expressing IFNγ and GzB, as well as the increased tumor antigen-induced CD8 T cell proliferation, demonstrate that combining CD73 inhibition and Ad-TK/Ad-Flt3L gene therapy resulted in increased anti-tumor CD8 T cell effector functions.

## Discussion

A critical factor influencing the glioma TME is the mutation status of IDH1. IDH1-R132H (mIDH1) is a prevalent gain-of-function mutation leading to an accumulation of the oncometabolite 2-hydroxygluatrate (2HG)^6^. 2HG competitively inhibits alpha-ketoglutarate-dependent enzymes responsible for DNA and histone demethylation^7^. This inhibition leads to a genome wide hypermethylation state causing significant alterations in the biology of glioma cells and the glioma immune response^4,34^. Notably, mIDH1 has been shown to increase the anti-tumor immune response by decreasing glioma cell PD-L1 expression and increasing granulocyte-colony stimulating factor (G-CSF) secretion^10,11^. PD-L1 expression was shown to be decreased due to increased DNA hypermethylation at the *CD274*, the PDL1 encoding gene, promoter region^10^. Glioma cell G-CSF production increased due to increased *CSF3*, G-CSF encoding gene, promoter enrichment for histone mark H3K4me3, which is a histone mark associated with transcriptional activation^11^. Our study uncovers another epigenetically driven mechanism, by which, the mutation of IDH1 influences the glioma immune response, thereby potentially altering efficacy of an immunotherapy.

CD73 is a cell surface ecto-nucleotidase that contributes to immune regulation through its primary function of converting extracellular AMP to adenosine^16,35^. Adenosine, in turn, acts upon adenosine receptors expressed on various immune cells, including anti-tumoral CD8 T cells, leading to suppression of anti-tumor immunity. This immunosuppressive signaling fosters a tumor-permissive microenvironment, particularly in gliomas where immune infiltration is already limited.

Due to its role in immune regulation, numerous cancer-targeting clinical trials have employed CD73 inhibition, mostly in combination with radiotherapies, chemotherapies, and other immune checkpoint inhibitor therapies^36,37^. Clinical trials investigating the inhibition of CD73 have indicated acceptable tolerability and a favorable safety profile. However, the early clinical efficacy data are heterogeneous. While some trials have shown signs of disease control, others have not demonstrated a clear clinical benefit (clinicaltrials.gov: NCT02503774, NCT04672434, NCT03381274, NCT03616886)^38–41^. Despite this mixed efficacy data and the current absence of CD73-targeting clinical trials specifically in gliomas, a growing body of preclinical evidence strongly supports the potential of using CD73 inhibition as a promising combinatorial treatment strategy for glioma^16,42^. As such, this study gives critical insight on the use of CD73 inhibition as a glioma immunotherapy based on the mutation status of IDH1.

Through integrated analyses of patient datasets, genetically engineered mouse model-derived glioma cells, and patient glioma biopsy-derived cells, we have demonstrated that mIDH1 induces DNA hypermethylation of the *NT5E* gene region leading to CD73 downregulation. Inhibiting DNA methylation restored CD73 expression, affirming causality between the IDH1 driven hypermethylation and *NT5E* gene silencing. This finding builds upon the paradigm, by which, mIDH1 affects the glioma TME by epigenetically downregulating gene expression.

Functionally, the lower CD73 expression on mIDH1 glioma cells leads to decreased extracellular adenosine in the glioma conditioned media. This translates to the decreased capacity of mIDH1 glioma cells to convert AMP to adenosine in the glioma tumor microenvironment. Previous studies have demonstrated the dominant role of adenosine in facilitating immune escape in gliomas, particularly by suppressing cytotoxic CD8 T cell activity^42^. The mIDH1-induced decrease in adenosine suggests that mIDH1 gliomas have less adenosine-mediated immune suppression. As a result, wtIDH1 gliomas may be more suitable targets for CD73 inhibiting therapies.

Using genetically engineered mouse models, we investigated the effect of CD73 on glioma survival. The relationship between CD73 and glioma survival was found to be complex. Despite the CD73 expression in wtIDH1 gliomas, the genetic knockout of CD73 conferred no survival benefit in the models. It is highly likely that even with the loss of the adenosine-mediated immune suppression, other mechanisms of immune suppression, such as the lack of anti-glioma CD8 T cells in the glioma TME, overcome any potential survival benefits. Therefore, we combined the targeting of CD73 with immune-stimulating Ad-TK/Ad-Flt3L gene therapy. Ad-TK/Ad-Flt3L gene therapy, which has recently completed a phase I clinical trial (clinicaltrials.gov, NCT01811992), boosts the anti-tumor CD8 T cell immune response^43^. The CD73 inhibition works with Ad-TK/Ad-Flt3L gene therapy by downregulating the immune suppression mechanism, which would otherwise suppress the cytotoxic CD8 T cells. These results add to the growing body of literature showing the effectiveness of CD73 inhibition when used in combination with other immune-stimulating therapies^16,37^. Thus, our data shows the necessity of combinatorial approaches when targeting CD73 in the glioma context.

Collectively, these findings establish a novel epigenetic mechanism linking mIDH1 to the regulation of adenosine-mediated immunosuppression. Our work provides preclinical evidence demonstrating the strategy of combining CD73 inhibition with TK/Ad-Flt3L gene therapy to improve patient prognosis in wtIDH1 glioma. These insights suggest the use of CD73 inhibiting therapies as a precision therapy for wtIDH1 glioma patients. This study also (1) supports considering IDH1 mutation status as a variable when planning glioma treatment regimens and (2) provides a basis for the of use CD73 inhibition as part of a combinatorial therapy for solid tumors marked with many immune suppressive mechanisms as adenosine reduction alone may not be strong enough to combat tumor progression.

## Materials and Methods

### Animal Use

All mouse experiments were performed in accordance with the University of Michigan’s Institutional Animal Care and Use Committee. All animals were housed in an AAALAC accredited animal facility. The adult mice used were 8-12 week old C57BL/6 mice (Jackson Laboratory, 000664). For generating CD73-KO glioma models, Nt5e^tm1Lft^/J mice (Jackson Laboratory, 018986) were used.

### Genetically Engineered Murine Glioma Models via the Sleeping Beauty Transposon System

Murine models of CD73-KO gliomas were created using the Sleeping Beauty (SB) Transposon system^21^. Tumors then develop intracranially *de novo* with the chosen genetic lesions. The genetic lesion encoding plasmids include: SB transposase and luciferase (pT2C-LucPGK-SB100X; SB-Luc), a short hairpin against p53 (pT2-shp53-GFP4; shp53), a short hairpin against ATRX (pT2-shATRX53-GFP4; shATRX), and a NRAS-G12V activating mutation (pT2CAG-NRASV12; NRAS), with or without the plasmid mutant IDH1-R132H (pKT-IDH1(R132H)-IRES-Katushka; mIDH1)^11^. Plasmids can be found on AddGene (Table S1) The genotype of the genetically engineered tumor cells have these combinations: shp53, shATRX, NRAS, Nt5e^tm1Lft^, (wtIDH1, CD73-KO) and shp53, shATRX, NRAS, Nt5e^tm1Lft^, IDH1-R132H (mIDH1, CD73-KO). The SB protocol was performed as previously described^11^. In short, plasmids were mixed (20 µg plasmid in a total of 40 µL plasmid mixture) with *in vivo*-jetPEI reagent (Polyplus Transfection) (2.8 µL per 40 µL plasmid mixture) and glucose (10%). The transfection mixture was incubated for 20 minutes at room temperature and then moved on ice. The lateral ventricle (1.5 mm rostral and 0.8 mm lateral from lambda; 1.5 mm deep) of neonatal Nt5e^tm1Lft^/J mice was injected with 0.75 µL plasmid solution at a rate of 0.5 µL/min. Over weeks, the tumors develop de novo.

### Generation of Primary Mouse Glioma Cells

Mouse glioma cells were generated by harvesting brain tumors at the time of euthanasia. The brains were removed, and tumors were identified/excised by GFP expression under an epifluorescence microscope. The tumor mass was cut into small pieces and disassociated chemically using StemPro Accutase (Thermo Fisher Scientific, A1110501), filtered through a 70 µm strainer and cultured in cell medium [DMEM/F12 with L-Glutamine (Thermo Fisher Scientific, 11-320-033), B-27 supplement (1X) (Thermo Fisher Scientific, 12-587-010), N-2 supplement (1X) (Thermo Fisher Scientific, 17-502-048), Antibiotic-Antimycotic (1X) (Thermo Fisher Scientific, 15-240-062), and Normocin (0.1 mg/mL) (InvivoGen, ant-nr-1)] at 37 °C, 5% CO_2_. FGF and EGF (Peprotech, 100-18B and AF-100-15, respectively) were added twice weekly at 1 µL (20 ng/µL each) per 1 mL medium. Cells expressing GFP and Katushka were sorted by FACS.

### Intracranial Implantations

The syngeneic glioma mouse models were prepared by implanting ketamine (75 mg/kg) and dexmedetomidine (0.5 mg/kg) anesthetized mice with 30,000 NRAS-G12V/shATRX/shP53±IDH1-R132H harboring glioma cells in the right striatum. The implantation was conducted with a 30-gauge Hamilton syringe on a stereotactic fixation device (Stoelting). The coordinates for implantation are 1.00 mm anterior and 2.00 mm lateral from the bregma and 3.25 mm ventral deep from the dura. Glioma cells were injected at a rate of 1 μL/min. Atipamezole (1 mg/kg) was administered as the sedative reversal agent. Mice were given buprenorphine (0.1 mg/kg) and carprofen (5 mg/kg) for analgesia.

### Cell Culture

All cells were cultured at 37 °C (5% CO_2_). Glioma cells NPA, NPAI, NPA-OVA, NPA-CD73KO were grown in DMEM/F12 medium (Thermo Fisher Scientific, 11320033) supplemented with L-Glutamine (5mM; Thermo Fisher Scientific, 25030081), B-27 supplement (1X; Thermo Fisher Scientific, 14190144), N-2 supplement (1X; Thermo Fisher Scientific, 17502048), Antibiotic-Antimycotic (1X; Thermo Fisher Scientific, 15240062), and Normocin (0.1mg/mL; InvivoGen, ant-nr-1). FGF (PeproTech, cat. no 100-18B) and EGF (PeproTech, cat. no. AF-100-15) were added twice weekly at 1 µL (20 ng/µL each stock, 1000X formulation) per 1 mL medium. RPA, RPAI, and SF10602s were grown in Neurobasal medium (Thermo Fisher Scientific, 21103049) supplemented with L-Glutamine (5mM; Thermo Fisher Scientific, 25030081), B-27 supplement minus vitamin A (1X; Thermo Fisher Scientific, 12587010), N-2 supplement (1X; Thermo Fisher Scientific, 17502048), Antibiotic-Antimycotic (1X; Thermo Fisher Scientific, 15240062), and Normocin (0.1mg/mL; InvivoGen, ant-nr-1). FGF (PeproTech, cat. no 100-18B) and EGF (PeproTech, cat. no. AF-100-15) were added twice weekly at 3 µL (20 ng/µL each stock, 1000X formulation) per 1 mL medium. SJGBM2-wtIDH1 and SJGBM2-mIDH1 glioma cells were grown in IMDM medium (Thermo Fisher Scientific, 12440061) supplemented with 10% heat inactivated fetal bovine serum (FBS; Thermo Fisher Scientific, A5256701). MGG8-wtIDH1 and MGG8-mIDH1 glioma cells were grown in DMEM medium (Thermo Fisher Scientific, 12430062) supplemented with 10% heat inactivated FBS. The MGG8-mIDH1 glioma cell culture media also included 100 units/mL puromycin (Gold Biotechnology, P-600-1) as a selective agent.

### In vitro mutant IDH1 Inhibition

SF10602s cells were seeded at a density of 1.0 × 10^6^ cells into laminin (Thermo Fisher Scientific, 23017015) coated, 25 cm^2^ flasks containing the SF10602 cell medium. Vorasidenib (AG-881; Selleck Chemicals, S8611) was prepared in DMSO. SF10602 cells were treated with a final concentration of 0.25 µM vorasidenib on days 1, 3, 5, 7 post seeding. On d8, the cells were collected to measure CD73 expression of CD8 T cells by flow cytometry.

### In vitro DNA Methyltransferase Inhibition

NPA and NPAI cells were seeded at a density of 5,000 cells into the wells of a laminin coated 6 well plate containing the NPA cell medium. Decitabine (DAC; Selleck Chemicals, S1200) was prepared in DMSO. NPAI cells were treated with a final concentration of 10 nM and 30 nM decitabine on days 1, 3, 5, 7 post seeding. On d8, the cells were collected to measure CD73 expression of CD8 T cells by flow cytometry.

### Tissue Processing to Single Cell Suspension for Analysis

Mice were monitored while the tumor develops and euthanized upon reaching the late-stage symptomatic phenotype. The tissues were processed as previously explained^44^. Briefly, the brains were removed, and tumors were identified/excised by GFP expression under an epifluorescence microscope. The tumor mass was cut into small pieces and disassociated chemically using StemPro Accutase (Thermo Fisher Scientific, A1110501) for 10 minutes; then it was pushed through a 70 µm cell strainer. The cell strainer was rinsed with two rounds of 10mL RPMI + 5% FBS to increase cell retrieval. To isolate the tumor-infiltrating immune cells, a Percoll (Fisher Scientific, 45-001-747) density gradient of 30% and 70% Percoll was employed. The tumor cell layer and immune cell layers were either analyzed same-day or frozen using freezing media [90% FBS:10% DMSO].

For splenocyte processing, freshly resected spleens were pushed through a 70 µm cell strainer. The cell strainer was rinsed with two rounds of 10 mL RPMI 1640 + 5% FBS to increase cell retrieval. For red blood cell (RBC) removal, the cells were centrifuged and resuspended in 1 mL 1X RBC lysis buffer (Biolegend, 420302) for 90 seconds. The lysis buffer was quenched with 9 mL PBS. The splenocytes were frozen using freezing media.

### Flow Cytometry

Cells were first stained with Live/Dead fixable stain (Thermo Fisher Scientific, L34966) for 20 minutes to label the dead cells. For antibody staining, cells were resuspended in sheath fluid (BD Biosciences, 342003), and the non-specific antibody binding was blocked with F_c_ block (anti-CD16/CD32). Cells were stained with fluorescence conjugated antibodies cocktail for 30 minutes at 4 °C. Post antibody staining, the cells were rinsed with sheath fluid. Flow samples were run on FACSAria II flow cytometer (BD Biosciences). The antibodies used for flow cytometry staining are listed in Table S2. Amcyan was used as a live/dead discrimination stain as only live cells are analyzed. Glioma cells in vivo were identified by Amcyan^-^CD45^-^GFP^+^. T cells were gated based on Amcyan^-^CD45^+^TCRb^+^CD8^+^. The expression, including the median fluorescent intensity, of molecules was calculated using FlowJo v10 (Becton, Dickinson and Company).

### Ad-TK and Ad-Flt3L Adenoviral Vector Production

Replication-deficient adenoviral vectors encoding Herpes simplex virus type 1 thymidine kinase (Ad-TK) and Fms-like tyrosine kinase 3 ligand (Ad-Flt3L) were produced in HEK293 cells. For each vector, sixteen 175 cm² flasks of HEK293 cells were cultured to 80–90 % confluency and infected at a multiplicity of infection (MOI) of 2. Infected cells were incubated until cytopathic effect was observed, after which viral particles were harvested by cell lysis. Crude viral lysates were subjected to two sequential cesium chloride (CsCl) gradient ultracentrifugation steps to achieve high-purity preparations, followed by three rounds of dialysis against Tris-buffered saline to remove residual CsCl. Purified viral stocks were aliquoted and stored at –80 °C until use. Viral titers were determined by endpoint dilution assay on HEK293 cells. Serial dilutions were plated in 96-well format, and cytopathic effect was assessed after seven days. Both Ad-TK and Ad-Flt3L reached final concentrations of approximately 1.3 × 10¹² PFU/mL, confirming efficient amplification and purification.

### Ad-TK/Ad-Flt3L Gene Therapy

Ad-TK/Ad-Flt3L gene therapy was started day 5 post intracranial implantations by the injections of adenoviral vectors at the site of glioma cell implantation. Mice were anesthetized with ketamine (75 mg/kg) and dexmedetomidine (0.5 mg/kg). Using a 30-gauge Hamilton syringe and a stereotactic fixation device to hold the mouse skulls in a fixed position, a 1.5 µL solution of 2.5 × 10^8^ pfu of Ad-Flt3L and 2.5 × 10^8^ pfu of Ad-TK was administered at a depth of 3.75 mm, 3.5 mm, and 3.25 mm (0.5 µL each location) from the dura. Ganciclovir (GCV; 7 mg/kg; MedChemExpress, HY-13637) was administered *i.p*. twice daily as displayed on the glioma treatment timeline, days 6 to 15 for the survival experiment and days 6 to 12 on the timed endpoint experiment.

### In vivo Glioma Monitoring

For *in vivo* imaging of glioma progression, bioluminescence was measured using an IVIS Spectrum Imaging System (Xenogen). For the *in vivo* imaging system (IVIS), the settings used were default except for the exposure time was set to automatic and field of view set to “D”. With the IVIS primed and settings configured, 100 µL of luciferin (30 mg/mL) solution was injected *i.p.*, and mice were then anesthetized with oxygen-isoflurane (2.5% isoflurane). 5 minutes post luciferin administration, the mice were placed in the IVIS and measured. To score luminescence, Living Image Software Version 4.3.1 (Caliper Life Sciences) was used with a circular region of interest (ROI) over the head. The luminescence intensity was measured using the calibrated units photons/s/cm^2^/sr. Other than IVIS monitoring, mice were monitored daily for signs of morbidity, including ataxia, impaired mobility, hunched posture, low responsiveness, and seizures.

### Analysis of TCGA Data

For *NT5E* mRNA expression, TCGA glioblastoma multiforme (GBM) and TCGA brain lower grade glioma (LGG) cohorts’ gene expression and IDH1 mutation status data were collected using cBioPortal and merged to form a GBMLGG merged cohort^45,46^. Glioma cases were stratified based on both IDH1 mutation status (wtIDH1 n=204, mIDH1 n=230) and compared based on the *NT5E* mRNA expression.

For the survival analysis, TCGA (GBMLGG) gene expression, clinical survival, and IDH mutation status data were analyzed using the GlioVis data visualization tool^47^. Glioma cases were stratified based on both IDH mutation status (wtIDH n=232, mIDH n=428) and *NT5E* 1^st^ and 4^th^ Quartile. 1^st^ quartile makes up *NT5E* low expression strata (wtIDH n=59, mIDH n=108). 4^th^ quartile make up *NT5E* high expression strata (wtIDH n=58, mIDH n=107).

The TCGA AML gene expression and IDH mutation status were downloaded from the cBioPortal^45,46^. AML cases were stratified based on IDH mutation status (wtIDH n=440, mIDH1 n=38).

## Real-Time Quantitative Polymerase Chain Reaction (RT-qPCR)

Two-step qRT-PCR was performed following the manufacturer’s instructions. Total RNA was isolated using the RNeasy Plus Mini Kit (Qiagen), and 2 μg of total RNA was reverse transcribed to complementary DNA with the AffinityScript QPCR cDNA Synthesis Kit (Agilent Technologies). Quantitative real-time PCR (qPCR) analysis was performed using the Fast SYBR™ Green Master Mix (Thermo-Fisher Scientific) in a ViiA™ 7 Real-Time PCR System (Thermo-Fisher Scientific). Mouse *NT5E* mRNA was amplified using primers 5′-GGGACATTTGACCTCGTCCA-3′ and 5′-CCACATGGATTCCACCCACT-3′, and human *NT5E* mRNA was amplified using primers 5′-AATGGTGGAGATGGGTTCCAG-3′ and 5′-GAGGGAGTCAGCAATACAGGG-3′. The relative expression level of *NT5E* mRNA was calculated using the 2^−ΔΔCt^ method, normalized to the housekeeping gene: 18S.

### AMP and Adenosine Measurement

Non-adherent glioma cells (NPA, NPAI, RPA, RPAI) were seeded at a density of 2.0 × 10^6^ cells into the wells of a 6-well plate in 2 mL of their respective culture medium (sans the phenol red in the medium). Adherent glioma cells (SJGBM2-wtIDH1, SJGBM2-mIDH1) were seeded at a density of 2.0 × 10^5^ cells into the wells of a 6-well plate in 2 mL of the SJGBM2 culture medium (sans the phenol red in the medium). All cells were cultured at 37 °C (5% CO_2_) for 48 hrs. The media was collected and centrifuged (800 × g) at 4 °C for 10 minutes. The media was then analyzed using the Adenosine Monophosphate Assay Kit (Cell Biolabs, MET-5160) and Adenosine Assay Kit (Cell Biolabs, MET-5090) in line with the manufacturer’s suggested protocol. The fluorometric detection was measured on the plate reader Tecan Spark with the excitation wavelength of 530 and emission wavelength of 590.

### Western Blots

Mouse and human glioma cells (1.0 × 10^6^ cells) were harvested, and total protein extracts were prepared in a RIPA lysis and extraction buffer (Thermo Fisher Scientific, Pierce, 89900) with 1X of Halt protease and phosphatase inhibitor cocktail (Thermo Fisher Scientific, 78442). 20 μg of protein extract (determined by bicinchoninic acid assay (BCA), Pierce, 23227) were separated by 4-12% SDS-PAGE (Thermo Fisher Scientific, NuPAGE, NP0322BOX) and transferred to nitrocellulose membranes (Bio-Rad, 1620112). The membrane was probed with 1:1000 of *NT5E*/CD73 (D7F9A) Rabbit mAb (Cell Signaling Technology #13160), 1:2000 of an anti-β-actin antibody (Cell Signaling Technology #4967); and secondary [Dako, Agilent Technologies, goat anti-rabbit 1:4000 (P0448), rabbit anti-mouse 1:4000 (P0260)]. Enhanced chemiluminescence reagents were used to detect the signals following the manufacturer’s instructions (SuperSignal West Femto, Thermo Fisher Scientific, 34095). WB quantification was performed using ImageJ. The CD73 band intensity was normalized to the β-actin housekeeping gene, and the reported data are from three replicates.

### Immunofluorescence

Mice were intracranially implanted with NPA glioma cells. 100 µL of 150 µg anti-mouse monoclonal antibody CD73 (TY/23; Bio X Cell, BE0209) was administered *i.p.* on days 7, 10, 13, 16 post glioma cell implantation. Mice were euthanized and perfused with Tyrode’s solution then fixed in 4% paraformaldehyde. Fixed brains were paraffin embedded and sectioned into 5 μm thick sections. Tissue sections were incubated with Alexa Fluor 546-conjugated anti-rat IgG (Thermo Fisher Scientific, A-11081) to detect tissue-bound circulating antibody (left panel). Nuclei were counterstained with DAPI (Thermo Fisher Scientific, 62247).

### Immunohistochemistry

The neuropathological evaluation of brains was done by first fixing them in 4% paraformaldehyde and embedding them in paraffin. 5 μm thick sections were obtained using (Leica RM2165) microtome system. Tissue sections were permeabilized with TBS-0.5% Triton-X (TBS-Tx) for 20 minutes, followed by antigen retrieval at 96 °C with 10 mM sodium citrate (pH 6) for 20 minutes.

The sections were allowed to cool to room temperature, followed by washing 5 times with TBS-Tx (5 minutes per wash), and blocked with 5% goat serum in TBS-Tx for 2 hours at room temperature. Brain sections were incubated with primary antibody GFAP (1:200), MBP (1:200), CD68 (1:1000), and Anti-Iba1 (1:2000) diluted in 1% goat serum TBS-Tx overnight at 4 °C. The following day, sections were again washed with TBS-Tx and incubated with biotinylated secondary antibody (1:1000 in 1% goat serum TBS-Tx) overnight at 4 °C. Biotin-labeled sections were subjected to 3, 3′- diaminobenzidine (DAB) (Biocare Medical) with nickel sulfate precipitation. The reaction was quenched with 10% sodium azide; sections were washed 3 times in 0.1 M sodium acetate, followed by dehydration in xylene, and cover-slipped with DePeX Mounting Medium (Electron Microscopy Sciences). Images at 10X and 40X magnification were obtained using brightfield microscopy (Olympus BX53).

### Hematoxylin and Eosin Staining

For histopathological assessment, paraffin-embedded 5 μm brain sections from each treatment group were stained with hematoxylin and eosin (H&E)^48^. Regions encompassing tumor tissue were imaged using the same microscope configuration. Liver tissues were processed similarly, sectioned at 5 μm, stained with H&E, and examined by bright-field microscopy (Olympus MA BX53).

### T cell Proliferation Assays

Mice were intracranially implanted with wtIDH1 glioma cells expressing ovalbumin (OVA). Mice were treated as described, and the spleens were processed and frozen. Splenocytes, in freezing media, were thawed in at 37° C for 1 minute before adding to RPMI 1640 medium. Splenocytes were then labeled with Carboxyfluorescein succinimidyl ester (CFSE; Fisher Scientific, 50-169-50). 500 µl of freshly prepared CSFE solution (5 μM) was added to the splenocytes, and cells were incubated for 5 minutes at 37 °C. The labeling was quenched by adding 5 mL of T cell media (1640 RPMI:10% FBS, 55µM BME, 2mM L-glut, 50µg/mL Pen-Strep, Normocin). The cells were incubated with the CFSE-T cell media solution for 5 minutes on ice to fully quench the CFSE. The cells were then centrifuged at 300 × g. The cells were washed 2 times with 10 mL of the T cell media solution, counted, and plated (96-well u-bottom) at a concentration of 100,000 cells in 200 μL of T cell medium. SIINFEKL (Anaspec, AS-60193-1) was added to the wells for a final concentration of 200 nM SIINFEKL. IL-2 (MilliPore Sigma, SRP3085) was added to the wells for a final concentration of 1 ng/mL. The plate was then transferred to and incubated at 37 °C, 5% CO_2_ for 4 days. Finally, the cells were stained for flow cytometry analysis of tumor cell specific CD8 T cell proliferation based on CFSE dilution. Tumor specific CD8 T cells were identified using PE-bound H2-K^b^ chicken OVA_257-264_ tetramer (courtesy of the NIH Tetramer Core Facility, NIH Contract 75N93020D00005 and RRID:SCR_026557).

### Methyl-seq

Genomic DNAs were isolated from mouse wtIDH1/mIDH1 NS by DNeasy Blood & Tissue Kit (Qiagen). DNA samples were processed at the University of Michigan Epigenomics Core for library preparation to measure 5-methylcytosine (5mC), 5-hydroxymethylcytosine (5hmC), and combined 5mC+5hmC marks. DNA concentrations were determined using the Qubit dsDNA quantitation kit, and quality was assessed with the Agilent 4200 TapeStation. For each library, 100 ng of genomic DNA was used. Libraries quantifying total 5mC+5hmC were prepared using NEBNext Enzymatic Methyl-seq (EM-Seq) Kit according to the manufacturer’s protocol. For specific 5hmC libraries, the EM-Seq protocol was modified as follows: stubby adaptors with incorporated 5hmC were used during ligation, and DNA underwent only beta-glycosylation prior to incubation with APOBEC, which was extended overnight. Final libraries were quantified by Qubit and examined for quality on the TapeStation using the HS D1000 DNA ScreenTape kit. Prepared libraries were pooled and sequenced on a NovaSeq 6000 S4 flow cell at the UM Advanced Genomics Core.

Read quality was assessed using FastQC (v0.11.8). Adapters and low-quality bases were trimmed with TrimGalore (v0.4.5) using the parameters: --adapter AGATCGGAAGAGC -e 0.1 --stringency 6 --length 20 --nextseq 20. Reads were aligned to the mm10 reference genome with Bismark (v0.22.1) using Bowtie2 (v2.3.4) under default settings (multi-seed length 20 bp, 0 mismatches). Duplicate reads were marked and removed using Picard (v2.20.2), and alignments with MAPQ <10 were filtered out using samtools (v1.2) with the parameter: -q 10. Methylation rates were called using MethylDackel (v0.4.0) with parameters -d 5 -D 2000 --mergeContext. Methylation mark deconvolution was performed using the MLML2R R package. Briefly, methylated and unmethylated counts were extracted from bisulfite-only and oxidative bisulfite-treated samples, and MLML2R::MLML() was applied to estimate genome-wide levels of methylcytosine (mC), hydroxymethylcytosine (hmC), and cytosine (C) using the package’s exact method.

### Serum Chemistry

Serum was collected in serum separation tubes (Fisher Scientific, 50-809-211). Samples in the serum separation tubes were left at room temperature for 45-60 minutes to allow blood coagulation before centrifugation at 2000 × g for 15 minutes at 4° C. Complete serum chemistry for each sample was determined by in vivo animal core at the University of Michigan by loading the serum in a Beckman Coulter AU480 (Beckman Coulter; Brea, CA) automated clinical chemistry analyzer.

### Statistical Analysis

All data analysis was performed using GraphPad Prism software (v10.4.1). Statistical significance was determined using the methods and sample size as described in the figure legends. Any p value of <0.05 was considered statistically significant.

### Data and Material Availability

Research reagents and data supporting the manuscript findings are available from the corresponding author M.G.C. upon request. All plasmids described in this study have been deposited in Addgene (Table S1).

## Acknowledgements

We thank Dr. J. Costello (University of California San Francisco) for providing the mIDH1 human glioma cells SF10602, which were a gift from the Dabbiere Family and funded by National Institutes of Health/National Cancer Institute (NIH/NCI) R01CA244838; the COG Repository at the Health Science Center for providing the human glioma cells SJGBM2; and Dr. J. Ohlfest (University of Minnesota, deceased) for providing the SB model plasmids. This work was supported by the NIH/National Institute of Neurological Disorder & Stroke (NINDS) Grants R37-NS144573-01, R01-NS122165, R01-NS122536, R01-NS124167, the NCI Cancer Center Support Grant 2P30CA46592, and the Rogel Cancer Center Faculty Scholar Award (to M.G.C.); NIH/NINDS Grants R01-NS122234 and R01-NS127378 (to P.R.L.); The Pediatric Brain Tumor Foundation, Leah’s Happy Hearts Foundation, Ian’s Friends Foundation, Chad Tough Foundation, and Smiles for Sophie Forever Foundation (to M.G.C. and P.R.L.); NCI Grants R01CA248160 and R01CA244931 (to C.A.L.). Rackham Merit Fellowship, Rackham Graduate Student Research Grant, Rackham Predoctoral Fellowship, Herman and Dorothy Miller Fund (to B.L.M.).

## Author Contributions

B.L.M., P.R.L., M.G.C. conceived and designed the study. B.L.M., J.A.P.A., A.A.M., A.A.D., Z.Z., S.R., M.L.V., C.T., K.B., A.W., C.C., L.H.Z., L.C.R., P.O., M.S.A., A.R., M.P., Y.W., B.S., P.S. conducted experiments and analyzed data. C.A.L., J.D.W., A.S., P.R.L, M.G.C. designed and supervised experiments. M.G.C. and P.R.L supervised the project. B.L.M and M.G.C. prepared the figures and wrote the article. All authors read and reviewed the article.

## Declaration of interests

Authors declare no competing interests.

## Notes

### Competing Interest Statement

The authors have declared no competing interest.

