## Supplemental Materials for "Reprogramming the Immune Suppressive Tumor Microenvironment in Glioma Enhances the Efficacy of Immune-Mediated Gene Therapy"

**Supplemental Material**

**Table S1: Plasmid Vectors**

| Plasmid number | Repository/Source | Plasmid Name | Insert |
|---|---|---|---|
| 124259 | Addgene | pT2-shATRX-GFP | shATRX |
| 124261 | Addgene | pT2-shP53 | shP53 |
| 124257 | Addgene | pKT-IDH1(R132H)-IRES-Katushka | R132H-IDH1 |
| 192869 | Addgene | pT2C-LucPGK-SB100X; SB-Luc | Luc |

**Table S2: Antibody and Cell Dye List**

| Antibody | Manufacturer | Cat # |
|---|---|---|
| *NT5E*/CD73 (D7F9A) Rabbit mAb | Cell Signaling Tech | 13160 |
| β-Actin Antibody | Cell Signaling Tech | 4967 |
| APC anti-human CD73 (Ecto-5'-nucleotidase) Antibody | Biolegend | 344006 |
| APC anti-mouse CD73 Antibody | Biolegend | 127210 |
| Carboxyfluorescein succinimidyl ester (CFSE) | Fisher Scientific | 50-169-50 |
| PE anti-mouse CD73 Antibody | Biolegend | 127206 |
| Alexa Fluor® 700 anti-mouse CD45 Antibody | Biolegend | 147716 |
| Live/Dead Fixable Aqua Dead Cell Stain Kit | Thermo Fisher Scientific | L34966 |
| Pacific Blue™ anti-mouse CD4 Antibody | Biolegend | 100531 |
| PerCP/Cyanine5.5 anti-mouse TCR-β chain Antibody | Biolegend | 109228 |
| Purified anti-mouse CD16/32 Antibody (Fc block) | Biolegend | 101302 |
| PE-bound H2-$K^b$ chicken $OVA_{257-264}$ tetramer | NIH Tetramer Core Facility | N/A |

| | | |
|---|---|---|
| PE/Dazzle™ 594 anti-mouse CD279 (PD-1) Antibody | Biolegend | 135228 |
| APC anti-mouse CD8a Antibody | Biolegend | 162306 |
| APC/Cyanine7 anti-mouse CD8a Recombinant Antibody | Biolegend | 155016 |
| Anti-Glial Fibrillary Acidic Protein (GFAP) Antibody | Millipore Sigma | AB5541 |
| Anti-Myelin Basic Protein (MBP) Antibody | Millipore Sigma | MAB386 |
| Anti-CD68 antibody | Abcam | ab125212 |
| Anti-Iba1 Antibody | Abcam | ab178846 |

Figure S1

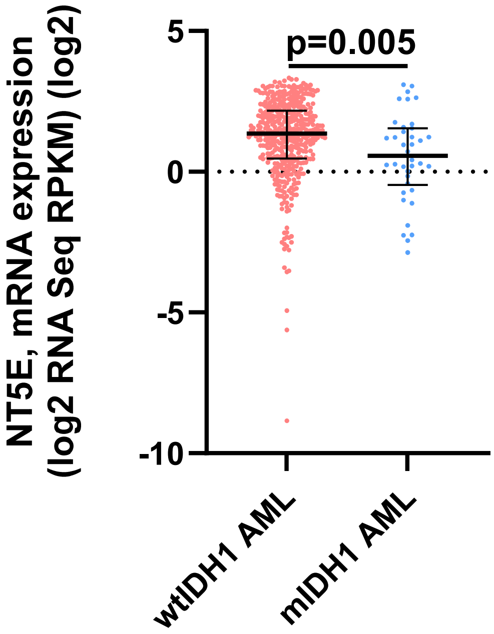

NT5E, mRNA expression
(log2 RNA Seq RPKM) (log2)
5
0
-5
-10
p=0.005
wtIDH1 AML
mIDH1 AML

**Figure S1: TCGA Analysis of CD73 Gene Expression in AML Patients Based on IDH1 Mutation Status**

Analysis of *NT5E* (CD73) gene expression wtIDH ($n$ = 440) and mIDH1 ($n$ = 38) from TCGA AML patients. RNA-seq data were obtained from TCGA (cBioPortal). Graph displays log2 expression value of *NT5E* mRNA expression (Mann-Whitney U test).

Figure S2

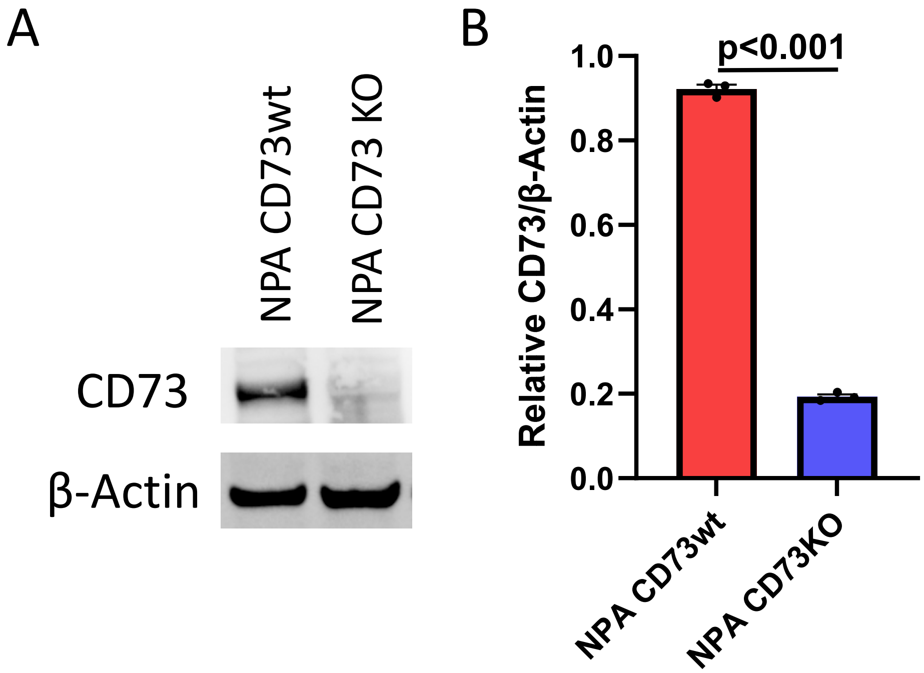
A
NPA CD73wt
NPA CD73 KO
CD73
β-Actin
B
p<0.001
1.0
0.8
0.6
0.4
0.2
0.0
Relative CD73/β-Actin
NPA CD73wt
NPA CD73KO

**Figure S2: Validation of the CD73KO in Mouse Glioma Cells**

(A) Western blot image and (B) quantitation of CD73 levels normalized to β-actin from NPA CD73wt and NPA CD73KO cell *in vitro* (technical replicates: $n$ = 3, Student's t test).

Figure S3

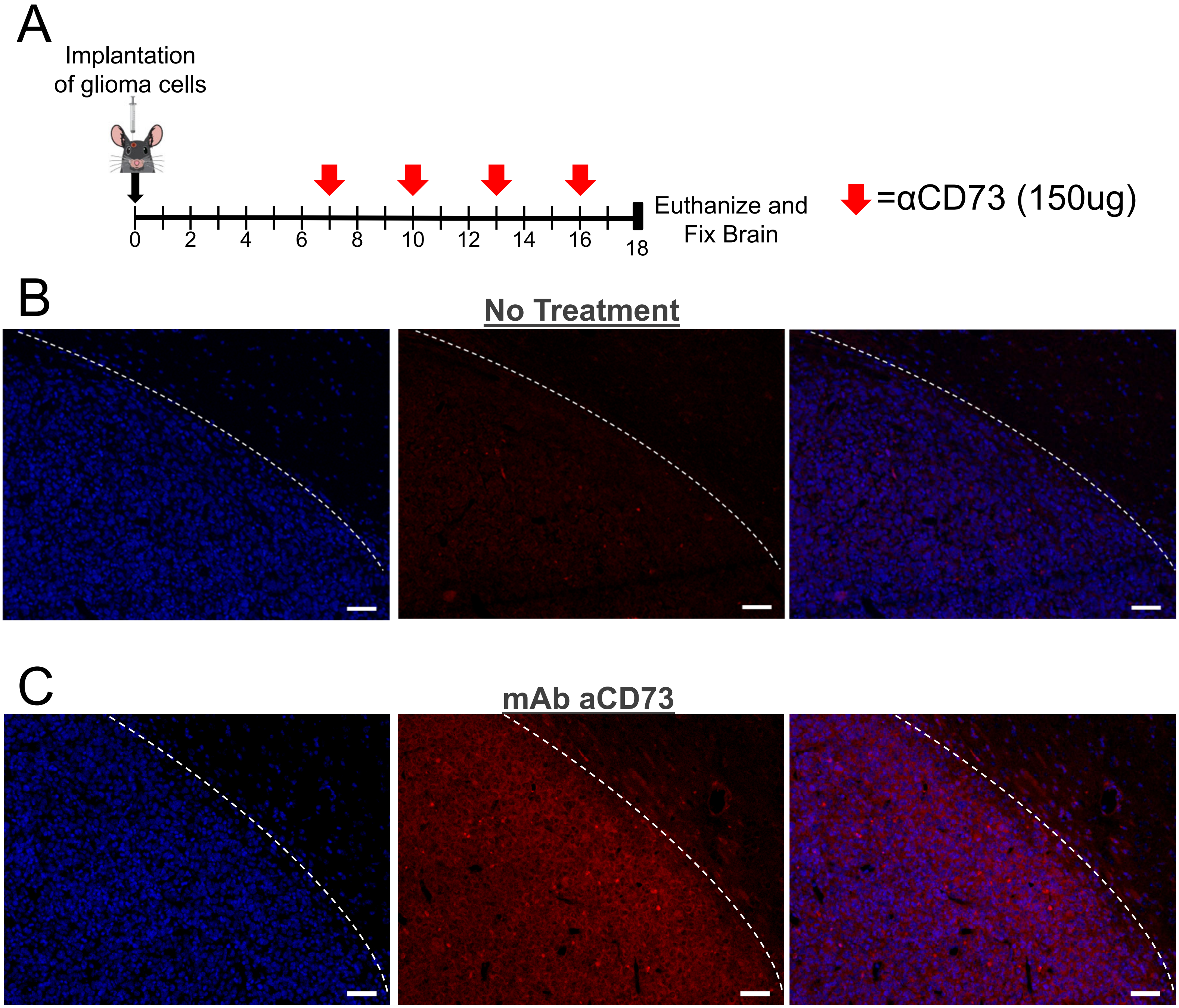


Scale bar: 50μm

**Figure S3: Antibody CD73 Inhibitor Successful Penetrates the Glioma Tumor**

(A) Diagram of glioma mouse model treatment timeline. Immunofluorescence analysis wtIDH1 glioma mouse models intraperitoneally systemically treated with an (A) isotype monoclonal antibody control (no treatment) or (B) CD73 monoclonal antibody (mAb aCD73). Tissue sections were incubated with Alexa Fluor 546-conjugated anti-rat IgG (red) to detect tissue-bound circulating antibody (center panel). Nuclei were counterstained with DAPI (blue; left panel). The DAPI and anti-rat IgG were overlayed (right panel) (scale bar, 50 μm).

Figure S4

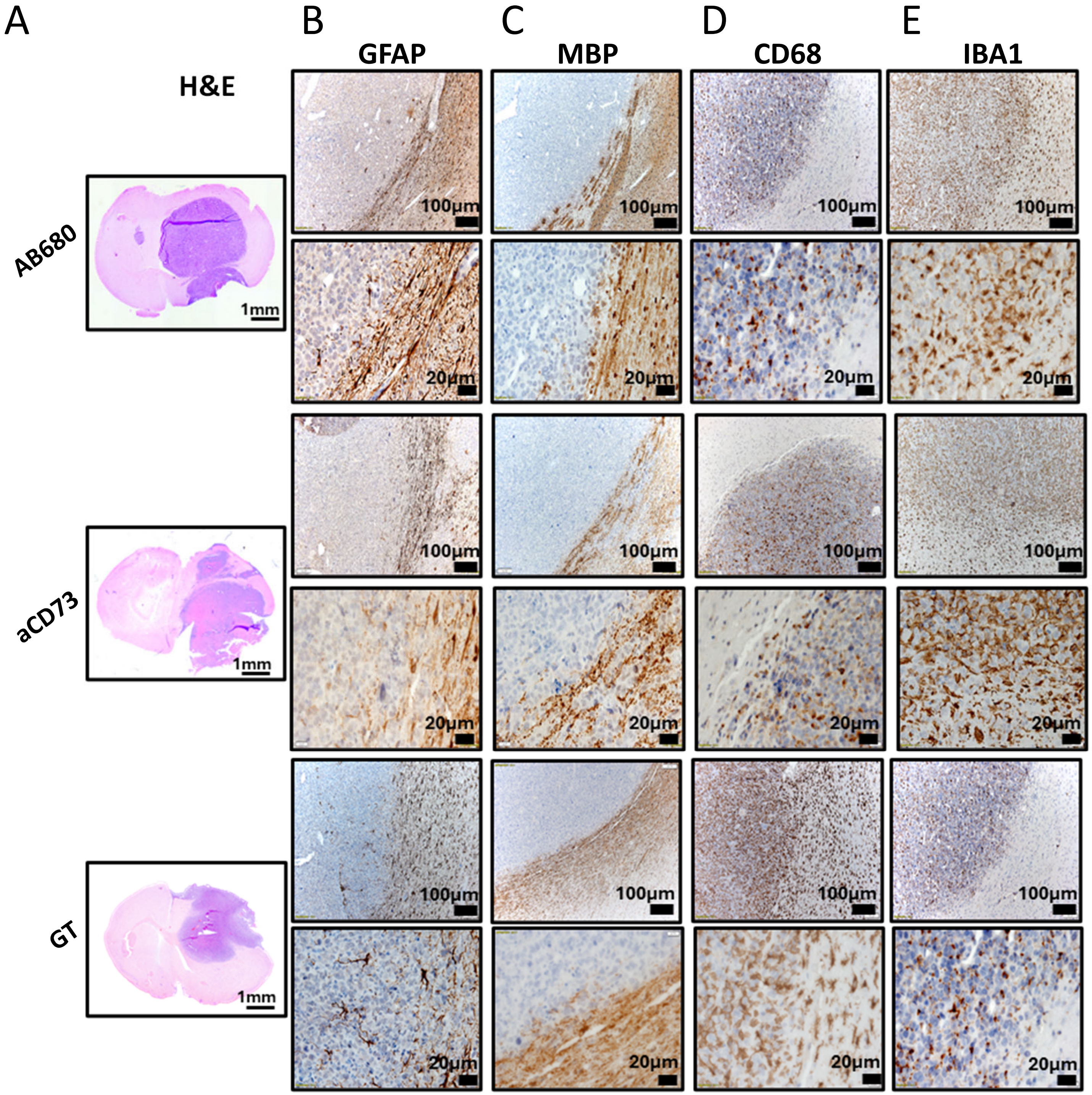

**Figure S4: Histopathological and Immunohistochemical Analysis of Brain Sections from Ad-TK/Ad-Flt3L Gene Therapy, Antibody CD73 Inhibitor, and AB680 in wtIDH1 Glioma Tumor-Bearing Mice**

(A) H&E staining of 5 µm paraffin-embedded coronal sections of brain from AB680, gene therapy, and CD73 antibody treatment groups (scale bar = 1 mm) shows tumor localization within the striatum of each group. Paraffin-embedded 5 µm brain sections for each treatment group were immunohistochemistry stained for glial and immune cell markers, including (B) glial fibrillary acidic protein (GFAP), (C) myelin basic protein (MBP), (D) CD68, and (E) IBA1. Low magnification lower panels (10X) panels (black scale bar = 100 µm). High magnification higher panels (40X) (black scale bar = 20 µm) indicate positive staining for the areas delineated in the low-magnification panels. Representative images from a single experiment, with independent biological replicates, are displayed.

Figure S5

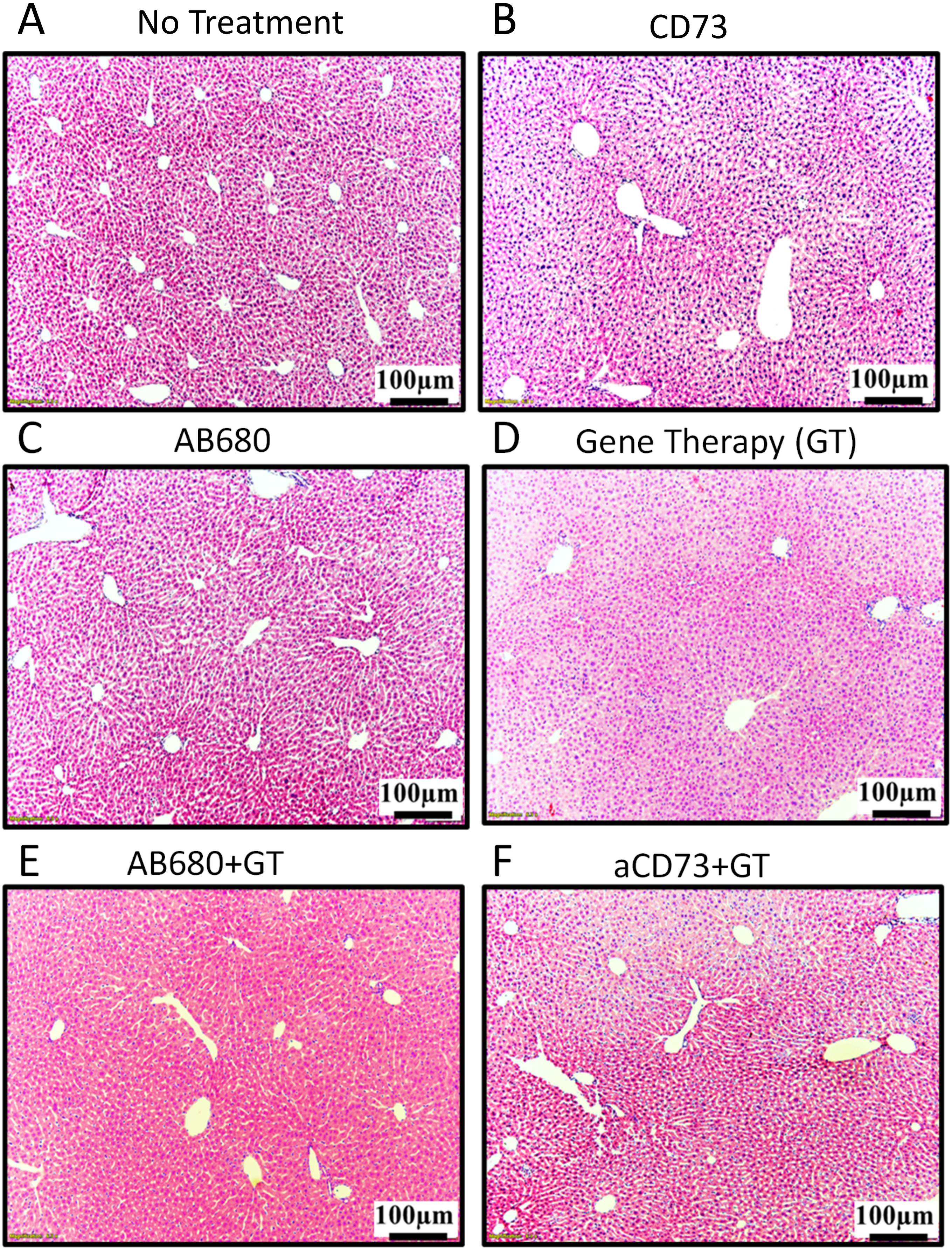

A
No Treatment
B
CD73
C
AB680
D
Gene Therapy (GT)
E
AB680+GT
F
aCD73+GT
100μm
100μm
100μm
100μm
100μm
100μm

**Figure S5: Histopathological Assessment of Livers from Tumor-Bearing Mice Treated with Gene Therapy and/or CD73 Inhibitors**

H&E staining of 5-micron paraffin-embedded liver sections from (A) no treatment control, (B) monoclonal antibody inhibitor of CD73 (aCD73), (C) AB680, (D) Ad-TK/Ad-Flt3L gene therapy (GT), (E) AB680 GT combination and (F) aCD73 GT combination treatment groups. Representative images from a single experiment, with independent biological replicates, are displayed (black scale bar = 100 μm).

### Figure S6

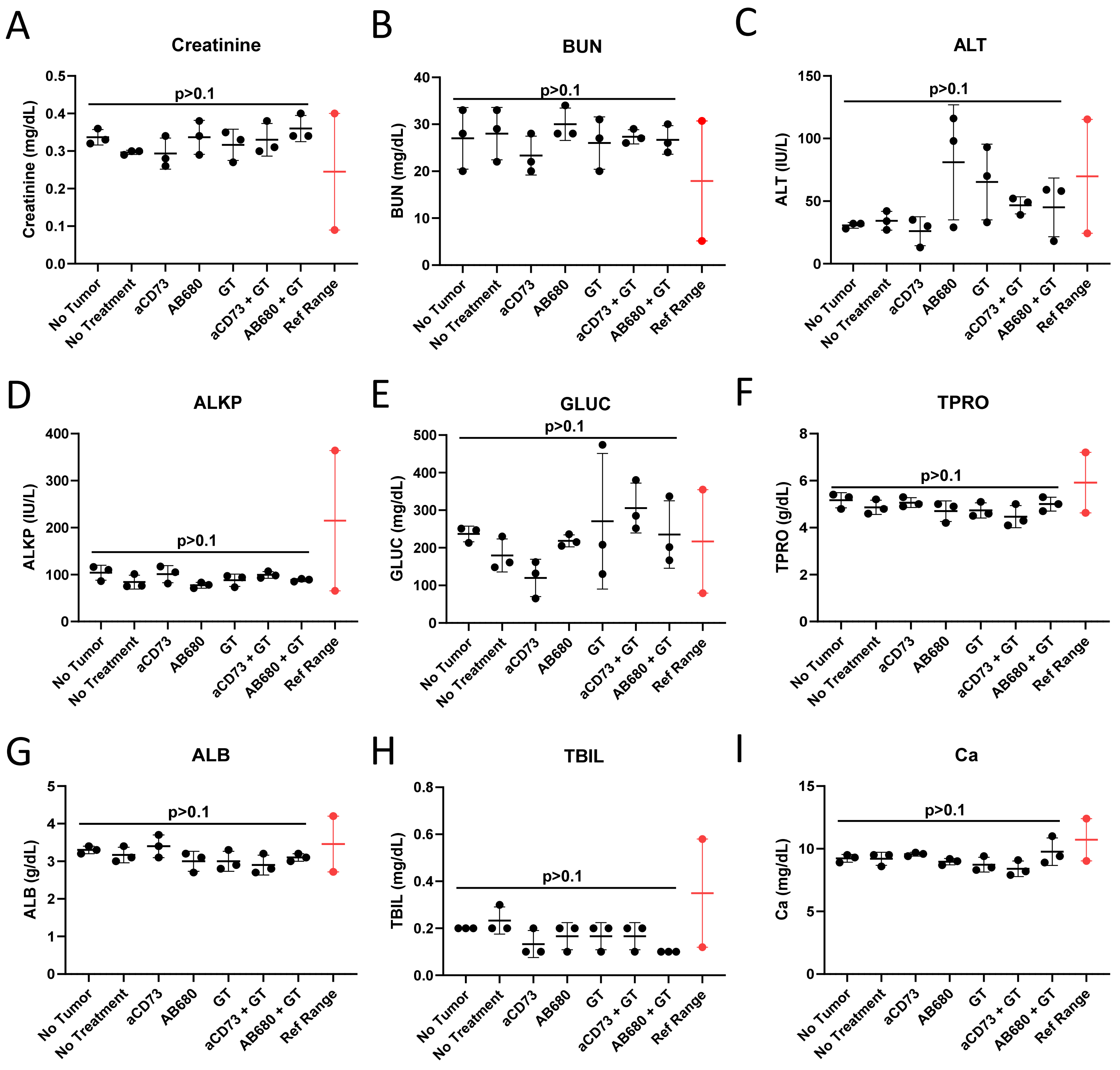
A
Creatinine
Creatinine (mg/dL)
p>0.1
B
BUN
BUN (mg/dL)
p>0.1
C
ALT
ALT (IU/L)
p>0.1
D
ALKP
ALKP (IU/L)
p>0.1
E
GLUC
GLUC (mg/dL)
p>0.1
F
TPRO
TPRO (g/dL)
p>0.1
G
ALB
ALB (g/dL)
p>0.1
H
TBIL
TBIL (mg/dL)
p>0.1
I
Ca
Ca (mg/dL)
p>0.1
No Tumor
No Treatment
aCD73
AB680
GT
aCD73 + GT
AB680 + GT
Ref Range

**Figure S6: Serum Chemistry of Tumor-Bearing Mice Treated with Gene Therapy and/or CD73 Inhibitors**

Mouse serum biochemical analysis following the conditions: no tumor, wtIDH1 glioma-bearing mouse models without treatment and treated with anti-CD73 monoclonal antibody (aCD73), AB680, Ad-TK/Ad-Flt3L gene therapy (GT), aCD73 GT combination, AB680 GT combination. The serum was collected at the late-stage symptomatic timepoint (just prior to euthanasia). For each condition 3 samples of serum were collected and the levels of (A) creatinine, (B) blood urea nitrogen (BUN), (C) alanine transaminase (ALT), (D) alkaline phosphatase (ALKP), (E) glucose (GLUC), (F) total protein (TPRO), (G) aluminium diboride (ALB2), (H) total bilirubin (TBIL), and (I) aalcium (Ca) were quantified (one-way ANOVA). The reference range is indicated in red.
